# Calcium triggers *Cryptococcus neoformans* aggregation by forming coordination bonds with capsular glucuronoxylomannan

**DOI:** 10.64898/2026.06.30.735596

**Authors:** Gracen Gerbig, Arturo Casadevall, Sindhoora Raja, Jason L. Sonnenberg, Maggie Wear

## Abstract

The cryptococcal polysaccharide capsule is a unique eukaryotic virulence factor that is a target for the immune system and the development of therapeutic antibodies. Our understanding of capsular architecture is limited to a few studies suggesting that metal dications play a role. In this work we explore a mechanism of cryptococcal aggregation that depends on calcium phosphate precipitation. We describe the chemical and biophysical properties of calcium interaction with the predominant cryptococcal polysaccharide, glucuronoxylomannan (GXM). We show that cell aggregation is a pH-dependent and occurs in a calcium dose-dependent manner. Furthermore, this cellular aggregation phenomenon as well as interpolymer capsular polysaccharide interactions are unique to calcium dications and do not occur with other mono- or dications as shown by size exclusion chromatography and circular dichroism. Diffusion ordered spectroscopy nuclear magnetic resonance and ab-initio calculations support complexation of calcium with glucuronic acid (GlcA). The ab-initio calculations also suggest that calcium ions can complex up to four GlcA monomers. Not only does calcium act as a scaffold for the cryptococcal capsule, interacting with up to four glucuronic acid residues of GXM, but calcium phosphate treatment of cells reduces the anti-phagocytic properties of the capsule, promoting ingestion by macrophages and altering antibody interactions with the capsule. This work advances our understanding of the cryptococcal capsule, its biophysical properties, by providing a model for the critical role of calcium interactions with capsular polymers of *Cryptococcus neoformans* including important impacts at the host-cell interface.

## Introduction

*Cryptococcus neoformans* is a yeast-like basidiomycete microorganism that poses a significant threat for immunocompromised individuals, being responsible for nearly 200,000 deaths worldwide each year, with a majority of cases occurring in people living with Human Immunodeficiency Virus (HIV) (1). Many deaths result from infection of the brain and meninges. Cryptococcus has a high neurotropism for the central nervous system, likely attributed to metalloproteinases and urease that promote blood-brain barrier entry and survival (2).

One of the most important and distinguishing virulence factors for *C. neoformans* is its polysaccharide capsule. *C. neoformans* changes its capsule in response to different stimuli, facilitating survival in different microenvironments. With a negative cell surface charge and lipophilic regions, the capsule may be able to modulate its composition or architecture to adjust repulsion or attraction to other cells and substratum, which affects virulence and survival of fungal cells (3, 4). The capsule is comprised of glucuronoxylomannan (GXM), accounting for approximately 90% of the non-water capsular mass, with the remainder being glucuronoxylomannogalactan (GXMgal), mannoproteins, and lipids (5). Capsular material is attached to the cell wall and becomes less dense and more porous as it extends outward (6, 7).

GXM is made of an *α*(1,3)-linked mannose (Man) backbone with *β*(1,2)- and *β*(1,4)-xylose (Xyl) and *β*(1,2)-glucuronic acid (GlcA) substitutions (8, 9). A polysaccharide with a mass of 1,700 – 7,000 kDa, it is the best studied component of the capsule (10, 11). A single GXM polymer may be comprised of repeating or a combination of different structural motifs, or triads. These triads vary according to the number and location of xylose residues (6, 12). While GXM repeat motifs vary in xylose content, each triad contains three Man and one GlcA residue. Additionally, the polymer may be O-acetylated along the Man backbone, which influences serological activity and affects the architecture and structure of the polysaccharide (13). Both GXM and GXMgal contain GlcA residues, which, at physiologic pH, are anionic due to the presence of a carboxylate moiety. These residues contribute to the overall negative charge of the capsule (3, 14).

Biophysical studies examining the structural properties of the capsule suggest a strong role for dication metals (M^2+^). Capsular growth and polysaccharide assembly rely upon, and self-aggregate in the presence of calcium (Ca^2+^) and magnesium (Mg^2+^), which interact with GlcA residues (15). One possible mechanism of self-aggregation would coordinate M^2+^ to the carboxylic acid/carboxylate (COOH/COO^-^) moiety of GlcA (15). According to the model proposed by Nimrichter and colleagues a 1:2 molar ratio of metal ions to GlcA residues in GXM is sufficient for cross-linking of polymers and ionic bridging, whereas higher concentrations of metal saturate the negative charges of GlcA and prevent bridging of GXM (15). Measurements of capsule elasticity at various Ca^2+^ concentrations showed that higher Ca^2+^ levels were associated with increased capsule rigidity, supporting the theory of polysaccharide cross-linking by M^2+^ (16). The role of dications in GXM ultrastructure are also implicated in the establishment of cryptococcal biofilms. These complex communities of aggregated cells require Ca^2+^ and Mg^2+^ for biofilm adherence and maturation; further, chelation of these metals slows growth and reduces GXM secretion into the biofilm matrix (17). Mg^2+^has also been observed to induce capsule growth of cells by enhancing the expression of capsule regulating genes (18).

Situated at the interface between the environment and the cell, the capsule of *C. neoformans* undergoes physical changes in response to chemical signals from abiotic and biotic sources, host cells interactions and tissue colonization (19, 20). Physical modifications to the capsule influence the cell’s adherence to different surfaces and likely influence cell surface hydrophobicity (CSH), or the affinity of a cell’s surface to a hydrophobic vs a hydrophilic substance (21). Though relatively understudied in *C. neoformans*, CSH can affect fungal virulence, biofilm formation, and is important in considering antifungal drug targets (21). There is some evidence that M^2+^ ions contribute to a change in CSH, thereby changing the cell’s adherence and dynamics, but we do not understand how this may affect *C. neoformans*.

In this study we explore cryptococcal aggregation at the molecular level, considering the role of metal dications in cell-cell interactions using theoretical and experimental approaches. At the atomic level, we focus on the negatively charged component of GXM, GlcA, and its ability to form multidentate structures with dication metals. Macroscopically, we investigated the parameters of Ca^2+^-mediated aggregation and capsular changes in response to M^2+^ binding. New data presented herein demonstrates that the metal dications associate with GlcA residues, visible with ATR-FTIR and NMR-DOSY spectroscopy. These interactions are supported by theoretical studies, where we find that these metals form energetically stable complexes with up to four GlcA residues. Our study results in two findings. First, we build onto a previously defined model of GlcA and M^2+^ coordination, whereby polysaccharide cross-linking and cellular aggregation results from covalent-type bonding between metal dications and oxygen atoms of glucuronic acid. Second, we propose a model for calcium-phosphate biomineralization of *C. neoformans* capsule, affecting capsular polysaccharide structure and macroscopic cellular characteristics. Together, our results demonstrate that Ca^2+^ can bind strongly and drive aggregative effects on the polysaccharide capsule of *C. neoformans*.

## Results

### *Cryptococcus neoformans* aggregates form in the presence of select metal salts

To track the effects of excess metal salts on cryptococcal cells we added 10 mM metal chloride salts (dissociated Ca^2+^ = 440-700 µM), a concentration within physiological conditions (1.2mM Ca^2+^)(22), to cryptococcal cells (5 x 10^5^) and observed their effects on cells by microscopy. Ca^2+^ and cadmium (Cd^2+^) but not magnesium (Mg^2+^) or strontium (Sr^2+^) salts produce cryptococcal cell aggregates that cannot be dispersed by vortexing (Figure 1). Cells aggregated by Ca^2+^ and Cd^2+^ formed raft-like aggregates, floating to the top of the wells. Cells treated with Mg^2+^ and Sr^2+^ remained unaggregated and settled to the bottom of the wells in the 24 well plate. Physical modifications to cells, such as heat-killing or sonication, did not preclude aggregation (data not shown). Treatment of cells with EDTA reversed the aggregation of *C. neoformans* (data not shown).

**Figure 1:**
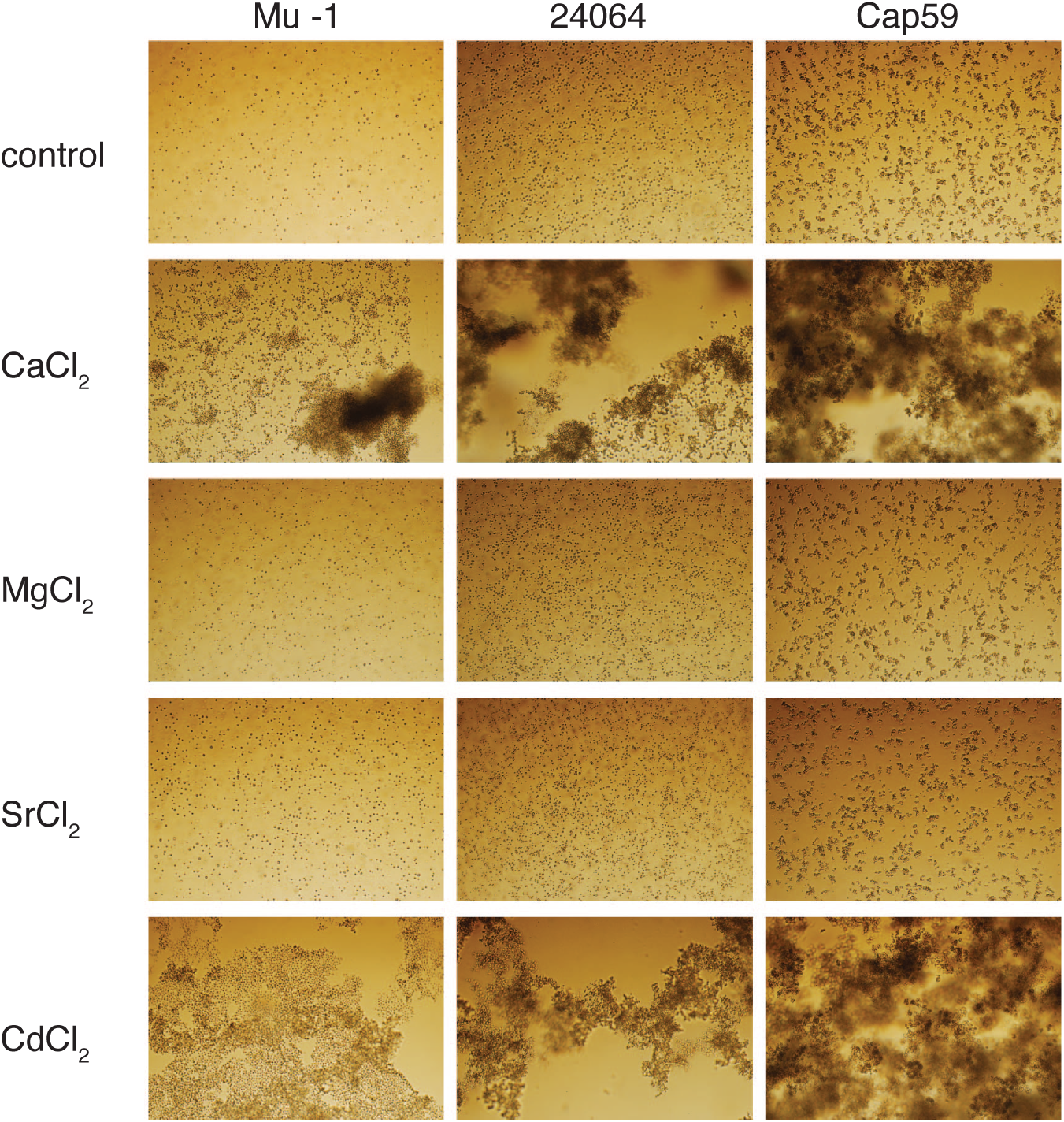
Cellular aggregation of *C. neoformans* cells in the presence of excess metal salts. Cells grown in YPD media for 48 h were seeded at 1x105 and 10 mM metal salts were added to a 24-well plate. Cryptococcal serotype A, GXM M2 motif expressing encapsulated strains, Mu-1 and ATCC 24064, along with acapsular mutant Cap59 (cap67) were used (left to right). Excess metal salts added (top down) control, CaCl2, MgCl2, SrCl2, CdCl2. (10X magnification).

### Dication Metal interactions with cryptococcal polysaccharide monosaccharides

The dominant cryptococcal capsular polysaccharide GXM has three different monosaccharides that could potentially interact with the metal dications to induce the observed aggregation, Man, Xyl, and GlcA. We performed diffusion ordered spectroscopy (DOSY) NMR to identify interactions between dication metals and the GXM monosaccharides. No significant diffusion differences were observed between Man and Xyl alone vs treated with dications (Supp. Fig. 1). But GlcA in the presence of M^2+^ showed different results in the DOSY experiments (Figure 2) compared to Man and Xyl. We then looked for interaction between metal dications and GlcA using Attenuated Total Reflectance (ATR) Fourier Transform Infrared (FTIR) Spectroscopy. ATR-FTIR spectrum analysis in the “fingerprint region” (1500-500 cm^-1^) (Supp. Fig. 2) shows a series of three peaks at 1070, 1416, and 1592 cm^-1^. The peak at 1592 cm^-1^ is typical of delocalized carboxylate stretching, and while this peak shifts with the addition of M^2+^, it is the broadening that suggests further stretching of the C-C bonds. The peak at 1070 cm^-1^ is indicative of C-O bond stretching, as may occur in an interaction between Ca^+2^ and the carboxylate of GlcA, and again this peak broadens with the addition of M^2+^. The peak at 1416 cm^-1^, on the other hand, indicative of C-H bond bending, does not broaden or change with the addition of M^2+^. Together these results suggest that stretching of C-C and C-O bonds occurs when M^2+^ is added to GlcA. The broadest peaks belong to Mg^2+^ and Ca^2+^, suggesting that they induce the most stretching of these bonds (Supp. Fig. 2) which led us to explore these interactions by DOSY NMR. These experiments showed that of three experimental groups, GlcA alone (Fig. 2A, 2B, blue), GlcA treated with calcium chloride (CaCl_2_) and phosphate (Fig 2A, red), or GlcA treated with MgCl_2_ and phosphate (Figure 2B, red) only calcium changed the diffusion constant. Ca^2+^ induces a broadening in the diffusion dimension corresponding to a change in diffusion constant from 5.11 x 10^-6^ to 5.49 x 10^-6^ cm^2^/ms, which is not observed with Mg^2+^. This change suggests an interaction between GlcA and Ca^2+^ but not with To better understand the molecular involvement between Ca^2+^ and glucuronic acid, DFT calculations were employed. Calculations modeling the glucuronic acid-metal dication complexes were used to identify the structures that are most likely to occur (23). The oxygen atoms in carboxylate groups have a delocalized, negative formal charge. Therefore, this is the most likely functional group to interact with metal dications. As such, three potential dication-GlcA coordination motifs, where GlcA acts as a bidentate ligand, were analyzed (Figure 3A). For coordination 3-A-1, the M^2+^ is positioned to coordinate exclusively with both carboxylate oxygens resulting in the formation of a four-member ring with the metal. In coordination 3-A-2, M^2+^ coordinates with one oxygen in the carboxylate group and the ring oxygen, forming a five-member ring. In coordination 3-A-3, M^2+^ coordinates to one of the oxygen atoms in the carboxylate group and ring carbon four hydroxyl, yielding a six-member ring. As six-membered rings are intrinsically more stable with lower energy arrangements than four-membered rings, conformer explorations using force fields showed that hydroxyl (-OH) groups bound to the pyranose ring achieved the lowest energy arrangement when all hydroxyl hydrogen atoms form intramolecular hydrogen bonds with a neighboring hydroxyl group oxygen atom. GlcA structures employed a so called “tail-chase” arrangement where all hydrogen bonds between pyranose ring hydroxyl groups pointed in the same direction. Such structures minimized the number of conformers resulting from hydrogen bonding between the hydroxyl (-OH) groups and focused the resulting reaction energies on the effects of metal-ligand bonding. These geometries were the initial starting structures for all DFT calculations on the complexes of Mg^2+^, Ca^2+^, Sr^2+^, or Cd^2+^ with GlcA ligands.

**Figure 2:**
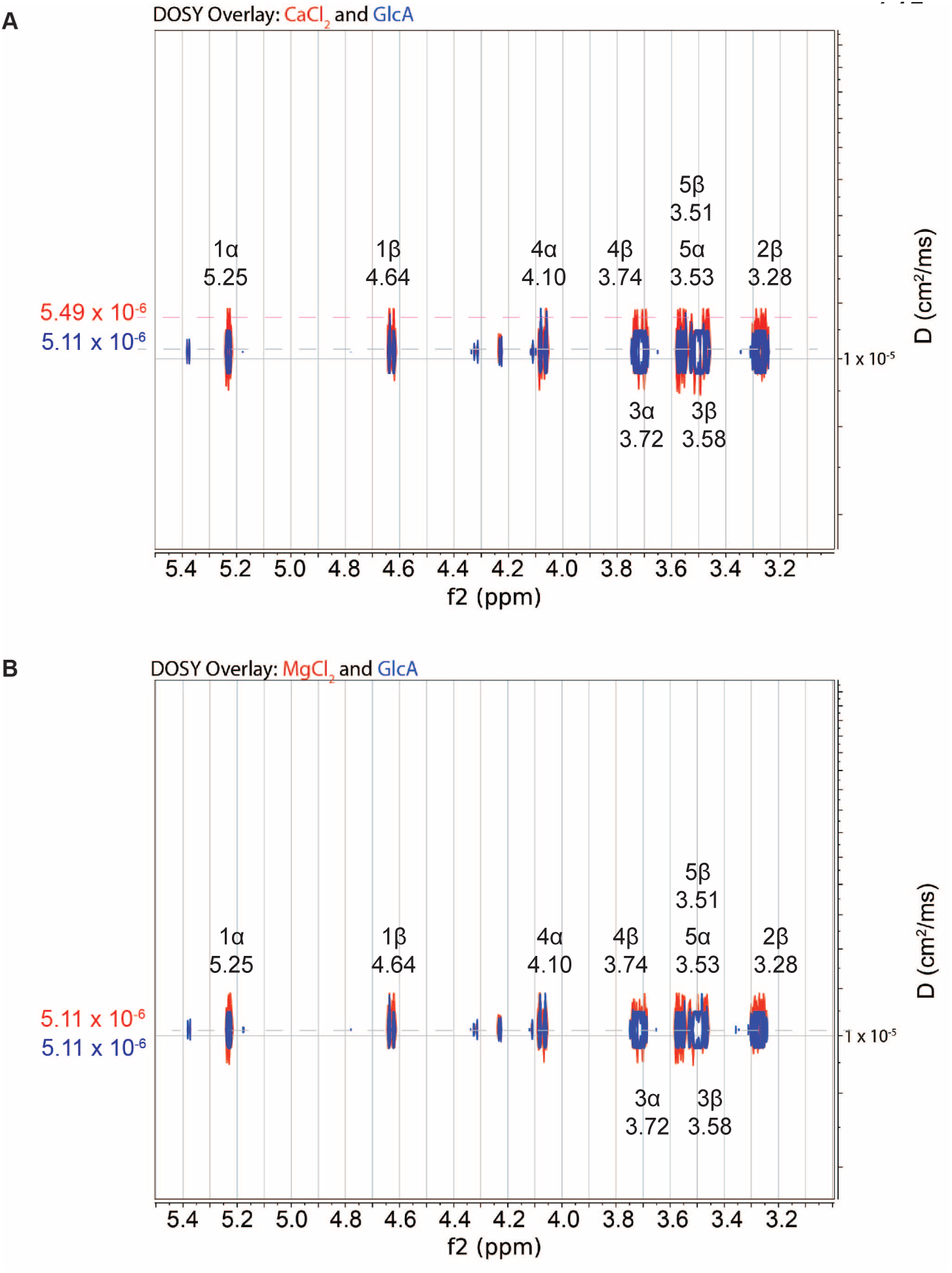
Diffusion NMR experiments to examine interaction between Glucuronic acid (GlcA) residues and calcium cations in the presence of phosphate. (A.) Top: NMR [1H] DOSY spectrum of GlcA in PBS (blue) overlayed with GlcA in PBS with calcium added (red). Bottom: NMR [1H] DOSY spectrum of GlcA in PBS (blue) overlayed with GlcA in PBS with magnesium added (red). Diffusion shifts noted on right-hand axis for each spectrum species overlay. (B.) Glucuronoxylomannan (GXM) motif repeats using glycobiology symbols.

**Figure 3:**
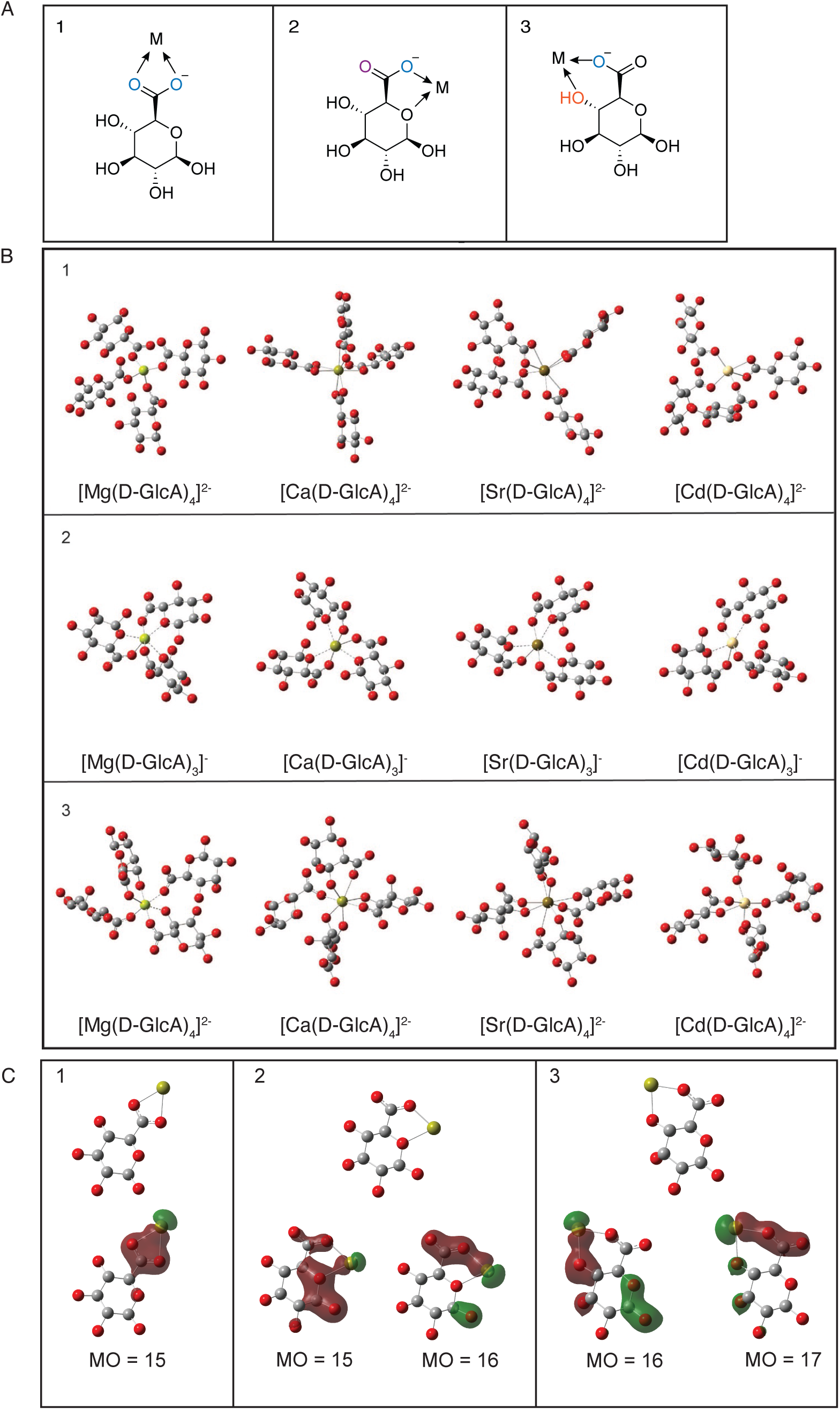
Density Functional Theory (DFT) modeling of interactions between GlcA and metal cations. A. Possible coordination events between metal (M) and GlcA oxygen residues based on negative charge density. 1. metal binding to carboxylic acid oxygens (blue); 2. metal binding to carboxylic acid oxygen (blue) and hydroxyl oxygen (orange); 3. metal binding to carboxylic acid oxygen (blue) and ring oxygen (purple). B. Optimized energetically favorable DFT structures of metal binding to GlcA residues based on above interaction schemes. Carbon atoms denoted in gray, oxygen denoted in red, magnesium in lime, calcium in yellow strontium in brown, and cadmium in white. Hydrogen atoms not shown. C. DFT modeling of molecular orbitals for calcium GlcA binding [Ca(GlcA)]. Selected molecular orbitals plotted of an isovalue at 0.035.

Enthalpies for individual structures were generated by computing the electronic energy and adding in the harmonic zero-point energy (Supp. Table 1). Motif 2 generates the lowest energy isomers for all four metals up to three GlcA ligands with the exception of magnesium bound to one GlcA where motif 3 is lower in energy by 0.28 kcal/mol. For calcium bound to one GlcA, motif 3 and motif 1 are 1.3 and 7.7 kcal/mol higher in energy respectively than motif 1, thereby emphasizing the energetic importance of ring formation. Reaction enthalpy was determined from the computed results for the following reaction where M^2+^ is the dictation metal and *n* is the number of GlcA in the reaction.

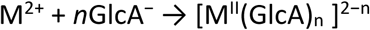

**Table 1:** Reaction Enthalpies (kcal/mol) for binding reactions and sequential addition of GlcA (Δ_r_H) M^2+^ + nGlcA- à [M^II^(GlcA)*_n_*]^2-n^ in solvato.

| Binding reactions | No. of GlcA | Mg | Ca | Sr | Cd |
| --- | --- | --- | --- | --- | --- |
| 1 (Figure 3-B-1) | 1 | -130.2 | -102.4 | -90.18 | -110.5 |
|  | 2 | -254.4 | -197.6 | -171.1 | -215.2 |
|  | 3 | -363.0 | -282.3 | -249.2 | -280.5 |
|  | 4 | -451.4 | -347.9 | -300.0 | -361.0 |
| 2 (Figure 3-B-2) | 1 | -171.9 | -134.5 | -115.2 | -122.9 |
|  | 2 | -366.5 | -274.5 | -242.5 | -253.3 |
|  | 3 | -436.5 | -352.7 | -308.0 | -299.9 |
| 3 (Figure 3-B-3) | 1 | -163.2 | -119.2 | -74.27 | -97.93 |
|  | 2 | -314.5 | -215.5 | -171.3 | -183.20 |
|  | 3 | -406.2 | -289.8 | -233.6 | -222.60 |
|  | 4 | -427.5 | -328.2 | -259.3 | -231.30 |

The reaction enthalpies (Δ_r_H) for the binding reactions of GlcA and M^2+^ were computed with incremental GlcA additions using an implicit solvent model (Table 1), showing the energetics of the three binding motifs proposed in Figure 3A. Metal complexation with the ring oxygen and the carboxyl group of GlcA was the most energetically stable with increasingly exothermic values at *n* = 1, 2 and 3 for all metals (Δ_r_H). Other coordination situations were also stable (negative in value), indicating exothermic reactions (Table 1). Additional analysis was performed to understand the result of incremental ligand binding on the average energy (or the change in reaction enthalpy across metal-oxygen bonds). Supporting Table 2 shows a pattern of decreasing exothermicity, consistent with increasing conformational constraints. Of the M^2+^-GlcA coordination modeling, the third coordination motif involving the carboxylate and ring oxygens (Figure 3A, third panel) had the most favorable average metal-oxygen energy for each metal. This is reflected in Table 1 where the enthalpic contributions of bonding of two GlcA ligands is *more than twice* as favorable compared to the reaction energy of bonding just one.

**Table 2.**
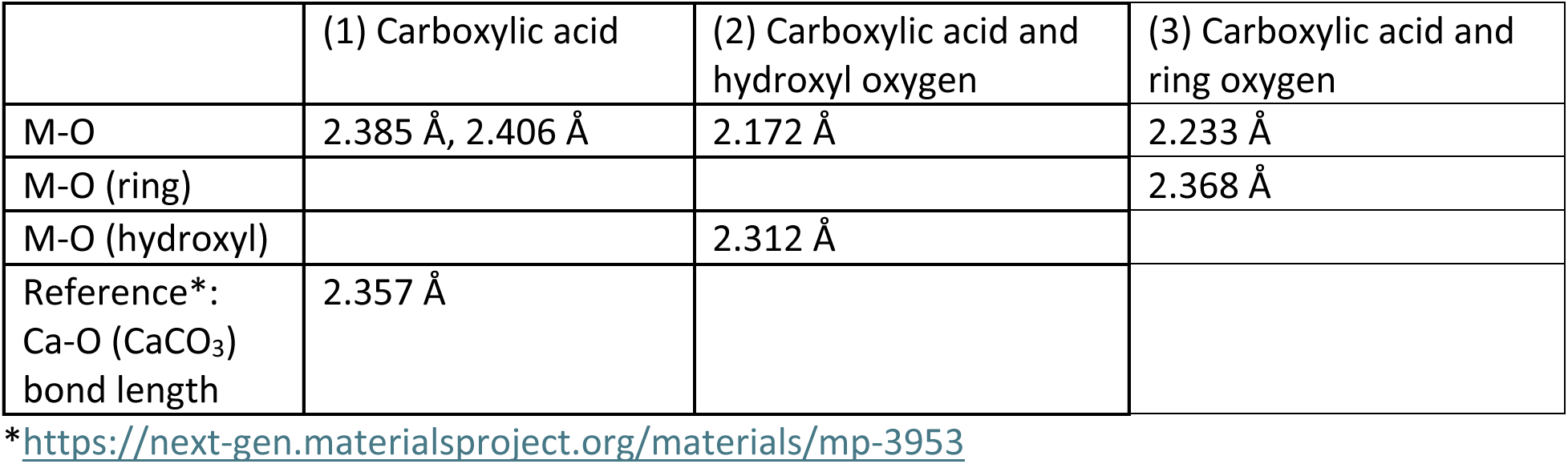
Bond lengths (Å) for calcium and glucuronic acid (n=1). Shaded boxes represent not applicable.

DFT-optimized structures are displayed in Figure 3B, where each panel shows the maximum ligand to metal ratio for each coordination motif. Figure 3B-1 shows the optimized structures for the metal coordination between the carboxylic acid oxygens of four GlcA molecules (n=4). Ligand coordination with metals can occur as monodentate, with a single binding site, bidentate, with two binding sites, or polydentate, with more than two binding sites, in this coordination reaction, Ca^2+^ binds more oxygens than Mg^2+^, Sr^2+^, and Cd^2+^, with four bidentate interactions. Mg^2+^ forms four monodentate interactions, Sr^2+^ forms three bidentate and one monodentate interaction, and Cd^2+^ forms one bidentate and three monodentate interactions. Figure 3B-2 shows the metal coordination of the metal situated between the carboxylic acid and a hydroxyl group attached to the 4^th^ carbon in the pyranose ring, giving two bidentate and two monodentate interactions for Mg^2+^, three bidentate and one monodentate interactions for Ca^2+^, four bidentate interactions for Sr^2+^, and two bidentate and two monodentate interactions for Cd^2+^. Geometry optimization by DFT calculations generally show Ca^2+^ coordinating more oxygens than the other metals, with Ca^2+^ preferring to have bidentate over monodentate interactions. Figure 3B-3 shows metal coordination between the pyranose ring and the carboxylic acid oxygens results in three bidentate interactions for each metal. In this coordination case, only three GlcAs can fit around the metals, but the reaction enthalpies of these products are more energetically favorable, having greater exothermic values when compared to the other coordination cases with the same number of GlcA.

### Covalent binding character in the case of calcium and glucuronic acid binding

To further understand the nature of the Ca^2+^ – oxygen bond, metal to oxygen bond lengths (Table 2) and bond angles (Table 3) were examined. Bond lengths can be predicted by adding the ionic radius of Ca^2+^ to the ionic radius of oxygen, and on average this bond is between 2.3-2.4 angstroms (Å) (24). M^2+^ – oxygen bond lengths for GlcA and Ca^2+^ dications are shortest for the coordination between Ca^2+^ and a carboxylate oxygen and a ring or hydroxyl group (Table 2). Shorter bond lengths indicate stronger bonds, due to increased electron density at the bonding interaction (25). The oxygen-metal-oxygen angles for these structures were the smallest for coordination at the carboxylate (1), followed by the carboxylate oxygen and the ring oxygen (3), and the carboxylate oxygen and the hydroxyl oxygen (2) (Table 3). Molecular orbitals (MOs) were visualized and analyzed by the percent orbital character. In each coordination mode, significant orbital overlap was observed between calcium and the oxygen atoms of GlcA (Figure 3C). Oxygen *s*-orbitals of GlcA donate electrons towards calcium’s empty 4*p* orbital, generating *σ*-bonding interactions. Analysis of bond lengths, bite angles, and molecular orbitals suggested that the most likely coordination motif for Ca^2+^ and GlcA was coordination at the carboxylate and pyranose ring oxygens (Figure 3C-3) > carboxylic acid oxygens (Figure 3C-2) > carboxylate oxygen and hydroxyl oxygen (Figure 3C-1).

**Table 3.** Bond angles (in degrees) for Ca^2+^ and GlcA.

|  | O-M<sup>2+</sup>-O angle |
| --- | --- |
| 1) Carboxylic acid | 56.6° |
| (2) Carboxylic acid and hydroxyl oxygen | 78.2° |
| 3) Carboxylic acid and ring oxygen | 69.1° |
| Reference*: Calcite (CaCO<sub>3</sub>) octahedral symmetry | 90 °, 180 ° |

\*https://next-gen.materialsproject.org/materials/mp-3953

### Effects of calcium on capsular polysaccharide

Size Exclusion Chromatography Multi-Angle Light Scattering (SEC-MALS), Matrix assisted laser desorption time-of-flight mass spectrometry (MALDI-TOF MS) and Circular Dichroism (CD) were performed to confirm a CaCl_2_ related change, as observed by DOSY NMR, in the capsular polysaccharide of *C. neoformans*. SEC-MALS analysis of our capsular polysaccharide preparation estimated the mass of polymers to be between 3.1 – 4.2 x 10^5^ Da and as large as 9.0 x 10^5^ Da, indicating at least two distinct populations (Figure 4A, top). Calcium or calcium-phosphate did not cause significant changes in estimated masses, but light scattering chromatograms showed significant differences in the elution times between the control CPS samples and the samples with calcium and calcium phosphate (Figure 4A). In addition to shifts in peak intensities and elution times, new peaks appear around 20 min that are absent in the control sample. Refractive index chromatograms also revealed differences in retention times between CPS samples. Compared to the CPS control, CPS treated with CaCl_2_ or with CaCl_2_ and phosphate showed earlier elution of peaks 1 and 2, while peak 3 eluted later (Figure 4A, bottom). MALDI-TOF MS analysis of the 20-minute SEC peak of 24064 CPS showed that the addition of CaCl_2_ yielded a single mass peak at 1238.64 m/z (calculated to be 2 Man + GlcA polymers and one Ca^2+^) while the addition of both CaCl_2_ and PO_4_ yielded a series of peaks including that at 1238.64 m/z. The largest of these peaks, 3221.87 m/z represents three GXM M2 polymers coordinated with a single calcium and phosphate (Supp. Fig. 3). Analysis of MS peaks were done using the GXM M2 motif component masses and represent the most likely, but not only, polymer combinations for the observed MS masses. When CPS was analyzed by Circular Dichroism (CD), only the CPS treated with CaCl_2_ manifested a change, with the CD spectra for CPS of strain 24064 shown in Figure 4B. While CPS control samples show a negative ellipticity, treatment with CPS and calcium resulted in a marked shift to a positive ellipticity. This shift is not observed when CPS is treated with calcium and phosphate. No shifts in ellipticity were observed for CPS treated with magnesium (data not shown).

**Figure 4:**
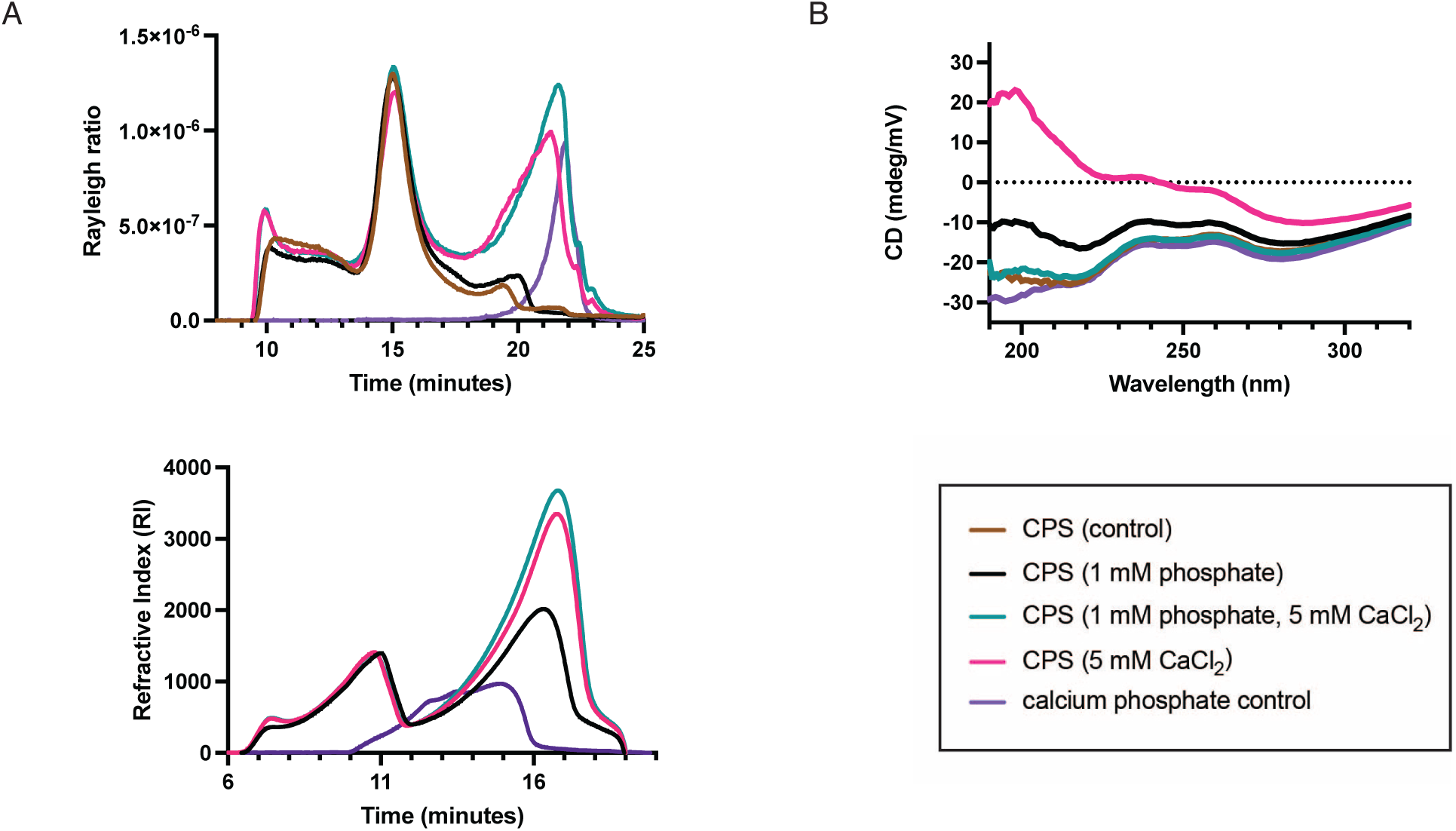
Experimental validation of DFT modeling using *C. neoformans* capsular polysaccharide (CPS). (A.) Light scattering and refractive index of Mu-1 CPS separated by size exclusion chromatography. <u>Top</u>: scattering intensity as defined by Rayleigh ratio, <u>Bottom</u>: Refractive index, visualization of polysaccharides. (B.) Circular dichroism spectra of CPS samples treated with CaCl2 or CaCl2 and phosphate. Brown and black lines compare CPS (control, no buffer) and CPS (1 mM phosphate) samples with no added dications. Pink and teal lines refer to CPS with 5 mM CaCl2 and CPS with 1 mM phosphate and 5 mM CaCl2 respectively. A calcium phosphate control (no CPS) is displayed

After examining the molecular interactions between dications and GlcA computationally and with isolated CPS, we returned to whole, encapsulated cells to further characterize dication metal-mediated aggregation. To determine the minimum amounts of calcium and phosphate required to produce cell-cell aggregation, we performed a calcium:phosphate titration (Figure 5). Though a minimum of 5 mM of CaCl_2_ and 1 mM phosphate produced aggregates of *C. neoformans,* cells aggregated in a concentration-dependent fashion with increasing concentrations of calcium and phosphate (Figure 5A). Scanning electron microscopy of *C. neoformans* cells treated with CaCl_2_ showed a more textured cell surface whereas the MgCl_2_ treated cells appear like untreated cells (Figure 5B). At an ultrastructural level, the microscopic CaCl_2_ aggregates appear amorphous and bridge neighboring cells.

**Figure 5:**
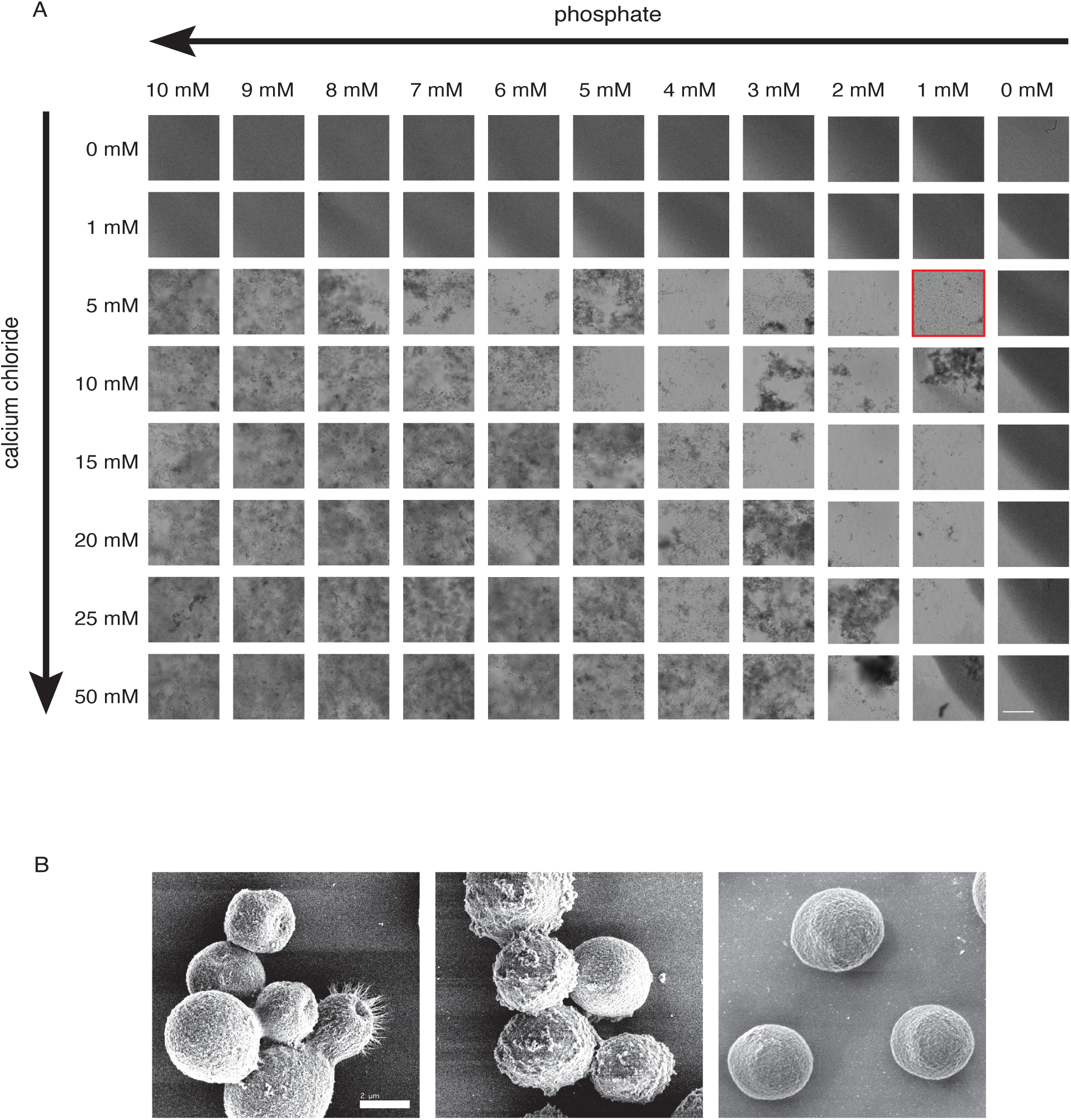
*C. neoformans* capsular interactions with calcium. A. Titration of CaCl2 and PO4 results in cellular aggregation with a dose-dependent response, increasing calcium and phosphate lead to increasing cell aggregation. First indication of aggregation noted with red square. (10x magnification, scale bar = 300 micrometers). B. Scanning Electron Microscopy (44) images of C. neoformans 24064 (left) 24064 in 1x PBS, (center) 24064 in 1xPBS + 10 mM CaCl2, (right) 24064 in 1xPBS + 10 mM MgCl2. (8500x magnification, scale bar = nm).

### Effects of calcium phosphate on phagocytosis

To test whether calcium phosphate induced aggregation affects phagocytosis, we compared the efficiency of Bone marrow derived macrophages (BMDMs) to phagocytose aggregated versus unaggregated cells. Antibody-mediated opsonization was used as a control, as the phagocytic efficiency of cells opsonized is significantly greater than that of unopsonized cryptococcal cells <u>(Casadevall et al. 1998)</u>. A modest increase in phagocytosis was noted between unopsonized and mAb-opsonized cells overall (Figure 6) (26). However, the phagocytic index (PI) of unopsonized cells treated with calcium phosphate was significantly higher than that of untreated, unopsonized *C. neoformans* (H99, P<0.05; Mu-1, P<0.0001; 24064, P<0.0001; Figure 6). Opsonized cells treated with calcium phosphate also showed elevated PI, though the effect varied by strain. While H99 showed no significant change, both Mu-1 (P<0.0001) and 24064 (P<0.001) exhibited significantly increased engulfment by macrophages following calcium phosphate treatment.

**Figure 6.**
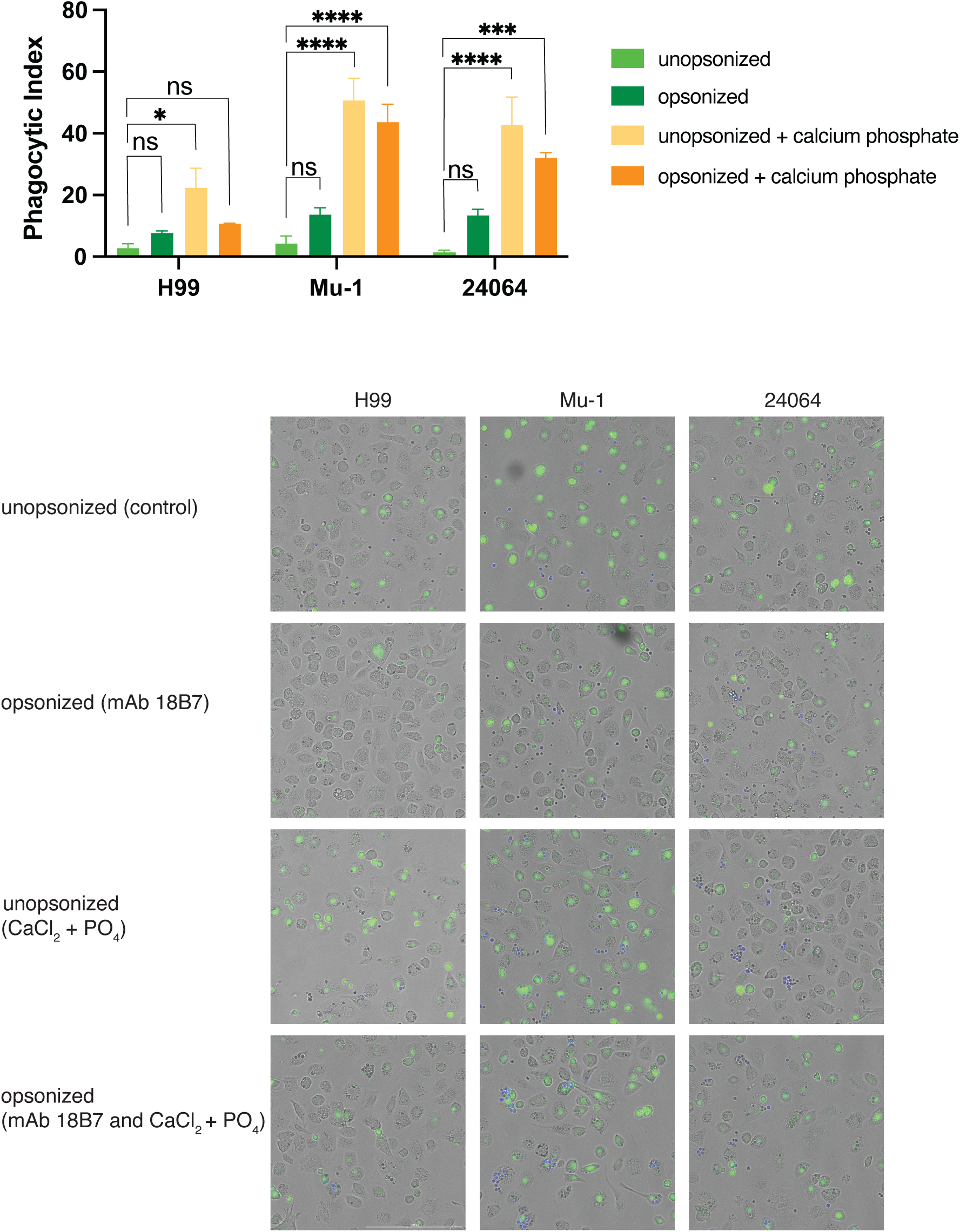
Phagocytosis of cryptococcal cells aggregated with calcium. (top) Calculated phagocytic index (bottom) representative images (10x magnification, scale bar = 300 micrometers).

Representative images highlight macrophages (stained green) with *C. neoformans* inside macrophages or outside (Figure 6). Internalized *C. neoformans* are clumped or single. This phenomenon is consistent across strains, with the phagocytosis index slightly increased in Mu-1 and 24064 as compared to WT H99. Supporting Figure 4 presents microscopy images of cells treated with both calcium phosphate and mAb 18B7. We observed that cells treated with calcium phosphate prior to mAb 18B7 staining exhibited stronger fluorescence and greater antibody binding than control cells. We also observed that the cells treated with calcium phosphate before mAb18B7 appeared aggregated and enmeshed by material (brightfield) and that the mAb 18B7 localized not only to the capsule surface (visible as a green ring surrounding the Uvitex 2B-stained cell wall) but also appeared to stain the surrounding matrix.

## Discussion

The major polysaccharide produced by the pathogenic fungi *Cryptococcus* is Glucuroxylomannan, or GXM, a branched polysaccharide consisting of a mannose backbone with xylose and glucuronic acid (GlcA) sidechains (8, 9). The negative charge of GlcA is thought to help buffer the pH of the environment for the cell (27). Previously, we reported that Ca^2+^ mediate self-assembly of GXM polymers to form the cryptococcal capsule (15). In this work we expanded on this observation examining how dications interact with the capsular polysaccharide of *Cryptococcus neoformans*. We found that the addition of positively charged Ca^2+^ in the presence of phosphate buffer initiated a cell-cell aggregation phenomenon reversible only upon chelation of Ca^2+^ ions. Of note, the concentrations of Ca^2+^ at which we observe induction of aggregation, when CaCl_2_ dissociation and absorption are taken into account (5 mM CaCl_2_ = 220-350 µM Ca^2+^) is below the bioavailable levels of extracellular Ca^2+^ (1.2 mM)(22, 28). This cell-cell aggregation observation holds across three different strains of *C. neoformans* despite varied capsular phenotypes. However, Ca^2+^ is not the only biologically relevant dication, nor the only dication with similar orbital geometry. Ca^2+^ along with Mg^2+^ and Sr^2+^ are group 2 alkaline earth (AE) metals and while Mg^2+^ forms interactions only with its valence electrons, the heavier AEs (*n*-1)*d* orbitals significantly affect binding causing them to behave more like transition metals (29). Therefore, we also included cadmium (Cd^2+^), a transition metal dication. The ionic radii for six coordinate complexes of Ca^2+^ and Cd^2+^ dications differ by 0.5 Angstroms (Å) whereas Sr^2+^ and Mg^2+^ differ by 0.18 and 0.25 Å respectively. Based on size alone, Ca^2+^ and Cd^2+^ should behave similarly, each forming up to six bonds. Ca^2+^ interactions with GXM are more similar to transition metals as NMR DOSY experiments show that the GXM residue interacting with dications is GlcA, and GlcA can interact with both Ca^2+^ and Cd^2+^, though to a lesser extent with Cd^2+^. While Cd^2+^ was able to induce cellular aggregation in *C. neoformans* strains, the association with GXM is limited, unlike that of Ca^2+^. This seems to reflect the biological availability of Ca^2+^ (4.4-7% of CaCl_2_ by mass) (28) and toxicity of Cd^2+^.

Previously, we modeled interactions between Mg^2+^ or Ca^2+^ and two GlcA residues of GXM in capsular assembly (15). Although Mg^2+^ and Ca^2+^ are both biologically available dications, they are structurally distinct and Ca^2+^, but not Mg^2+^, induced cell-cell aggregation. This is likely due to the difference in electron interactions wherein the Mg^2+^ valence shell is 3*s* and the Ca^2+^ valence shell is 4*s* leading to the involvement of the energetically accessible 3*d* orbitals. These 3*d* orbitals allow Ca^2+^, but not Mg^2+^, to interact with two GlcA residues as previously suggested, but also three, or even four GlcA molecules. DFT modeling showed that the polydentate GlcA residues could donate electron density to Ca^2+^. The most energetically favorable of these monodentate interactions was between Ca^2+^ the pyranose ring oxygen and the deprotonated carboxylic acid oxygen of GlcA. The oxygen-calcium bond is predominantly reported as ionic. On the Pauling scale, the difference in electronegativity between oxygen (3.5) and calcium (1.0) is large (2.5), supporting this characterization. Hard-soft Acid-Base (HSAB) theory posits that hard acids ted to interact with hard bases and soft acids tend to interact with soft bases. In other words, HSAB theory predicts that hard-hard interactions are more energetically favorable than hard-soft interactions. The dications Mg^2+^, Ca^2+^, Sr^2+^, and Cd^2+^ are hard acids while oxygen anions are hard bases, we expect the metals to ignore the other atoms in GlcA when oxygen anions are present (30, 26). The oxophillicity of metal cations is the basis for our modeling scheme. The -C(=O)OH group on the GlcA is deprotonated in our model giving the negatively charged -COO group three resonance structures: a delocalized one where the negative charge is spread evenly over both oxygens and two localized ones (-C(=O)O) containing a carbonyl group and a negative -O group. If a metal atom binds to both oxygen atoms than the delocalized resonance form predominates. If a metal atom binds to the deprotonated OH oxygen, the localized resonance form predominates. The data suggests that the GlcA:Ca^2+^ is not a non-specific physical interaction, as previously suggested, but rather has more covalent character (16, 31). Both bond length and bond angle affect interaction properties with ideal bond angles based on Valence-Shell Electron-Pair Repulsion (VSEPR) theory for Ca^2+^:O^-^ forming a pentagonal bipyramid or an octahedron coordination geometry being 60° in a bidentate interaction with an average bond length of 2.3-2.5 Å. The oxygen-metal-oxygen angle in GlcA:Ca^2+^ coordination closest to the ideal 60° is the carboxylate oxygen and the ring oxygen at 69.1° with an average bond length of 2.3 Å. While interaction with both carboxylate oxygens has a strained bond angle of 56.6° and average length of 2.4 Å and interaction with the carboxylate oxygen and the hydroxyl oxygen has a wide bond angle of 78.2° and average length of 2.24 Å. This bidentate bond angle and length modeling is corroborated by DFT modeling of molecular orbital overlap in the polydentate interaction as well as shifts in SEC retention time and CD molar ellipticity of cryptococcal CPS treated with calcium only and calcium and phosphate to compared to other dications and native CPS.

The GlcA:Ca^2+^ polydentate coordination mirrors that of EDTA and EGTA, which form stable coordinate covalent bonds (32). Forming a ring during coordination is a more energetically favorable process, with a lower energy state, than formation of an oxygen bond. Thus, we expected the Ca^2+^:GlcA coordination stability as follows, single bond to -COO^-^ group > pyranose oxygen + -COO^-^ group. In addition to VSEPR theory, Divalent Cation Bridging (DCB) theory, Alginate theory, and the Derajguin-Verwey-Landau-Overbeek (DVLO) theory have all been proposed to explain dication interactions with polysaccharides. Each theory highlights different aspects of particle interactions, with DCB focusing on ionic bridging, Alginate theory on the polysaccharide involvement, and DVLO on electrostatic/van der Waals forces. While DCB theory has been proposed as the mechanism by which self-aggregation of GXM occurs (15), the others have not been explored. DLVO describes the electric potential surrounding a particle, resulting in repulsion of neighboring particles, prohibiting aggregation, however, increased ionic strength of solution (like with additional Ca^2+^) compresses this, decreasing electrostatic repulsion and enabling van der Waals or other short-range attractive forces to drive aggregation (33, 34). Alginate theory, on the other hand, describes how cations interact with polysaccharides to create gels (33). The theory is specifically based on Alginate, a repeating mannuronic (ManA) and guluronic acid (GluA) polymer, wherein the gel is formed by calcium interactions with the negatively charged carboxylic acid of GluA resulting in an egg-box structure (33).

Our DFT calculations of GlcA coordination with dications showed that GlcA preferably forms bidentate interactions with Ca^2+^, and forms 4-6 heterocyclic rings depending on the location of oxygen coordination. Our initial molecular orbital (MO) calculations also show orbital overlap and mixing between the acidic oxygens of COO^-^, the pyranose ring oxygen, and hydroxyl oxygen attached to carbon 4. The ability of GlcA to bind dication metals with partial covalent character suggests a chelating-like role for the *C. neoformans* capsule. Further work will be needed to better clarify the polymer-level physical changes of whole GXM molecules induced by Ca^2+^. The egg-box structure of alginate gels, though mediated between calcium and GluA versus the GlcA of GXM, raises the possibility that calcium could mediate structured, cross-linked interactions between GXM motifs, not unlike those observed in the alginate model. A combination of these theories likely contributes to the calcium-induced aggregation we observe with *C. neoformans*; however, a key variable is missing from these models: phosphate. Our findings indicate that phosphate is a critical variable in the formation of cell clumps, but the ability of calcium to precipitate with anionic species in a manner that drives cell-cell association serves as a proof of concept that other biological buffers could similarly promote aggregation (35).

Knowing that GlcA:Ca^2+^ polydentate coordination forms bonds with covalent characteristics in the capsule and that the addition of surplus calcium phosphate results in cryptococcal cell aggregation, it is likely that this affects host-microbe interactions. The observed increase in phagocytic index of calcium phosphate mediated aggregated cryptococcal cells, versus unopsonized or GXM specific mAb 18B7 opsonized, suggests that calcium phosphate alters cryptococcal cell interaction with macrophages. Previously it was reported that cryptococcal aggregates and biofilms inhibit phagocytosis by macrophages (36, 37). This inhibition of phagocytosis was attributed to the overall size of cryptococcal biofilms along with the increased capsular size (36). The capsules of *C. neoformans* strain Mu-1, used in calcium phosphate mediated aggregation and phagocytosis experiments, are very large, yet we observed the opposite effect of aggregation on phagocytosis. Biofilms and cellular aggregation seem to have a protective effect on the survival of *C. neoformans* in other circumstances, but not when excess calcium phosphate is introduced. In addition to cellular aggregation, we observed microprecipitates in wells with excess calcium and phosphate suggesting that calcium-phosphate microprecipitates may not only form in the media, but on the surface of *C. neoformans* cells leading to biomineralization.

Microbial biomineralization involves the deposition of inorganic minerals and salts either within or around microbial cells. When Ca^2+^ is involved, the process is termed microbially induced Ca^2+^ precipitation (MICP) (38). Anionic residues on microbial surfaces serve as nucleation points for inorganic mineral formation. *Pseudomonas* spp. precipitate calcium in the form of apatite, amorphous calcium phosphate, or calcium carbonate by way of ureolytic and phosphate sequestering mechanisms (39). Urease positive fungi, such as *Neurospora* and *Pestalotiopsis* have been studied for their ability to biomineralize calcium and other dications (40, 41). The observation that urease-positive fungi also precipitate minerals may be explained by the carbonate anions generated during urea degradation, which can react with metals like calcium to form insoluble precipitates (41). *C. neoformans* is also a urease-positive yeast and hydrolyzes urea to ammonia to modulate the pH of its environment, particularly using it to survive the host phagolysosome (31, 42). A recent study investigating the interplay between urease production and melanization found that animals infected with urease-positive *C. neoformans* exhibited higher levels of ammonia gas and elevated tissue pH compared to animals infected with urease-deficient strains (43). These findings indicate that *C. neoformans* uses ammonia production to manipulate its microenvironment, potentially affecting neighboring cells. Given *C. neoformans’s* negatively charged capsule, with acidic GlcA residues capable of interacting with species in the environment, urease activity may also serve as a mechanism for promoting passive mineralization.

The hydrophilic nature of the capsule grants *C. neoformans* the ability to exist in an unaggregated state. Our DFT modeling indicates that Ca^2+^ polydentate coordination with GlcA provides a strong scaffold for the capsule. However, when excess Ca^2+^ is present this scaffold extends beyond the single cell, altering the cell surface and its hydrophobicity (CSH) to allow for increased cell-cell interactions, which may lead to cell aggregation. A measurement of a cell’s affinity for hydrophobic over hydrophilic surfaces, CSH greatly influences adherence to both abiotic and biotic surfaces. A recent study by our group found that CSH varies among *Cryptococcus* strains and serotypes, and proposed that differences in capsule composition, including variations in GXM motifs and side chain substitutions, might play a role in % CSH (44). This work also shows that the more hydrophobic strains are more easily preyed upon by amoeba, a natural predator of *Cryptococcus* in the environment. It has been proposed that phagocytosis by macrophages is mechanistically similar to that of amoeba predation (45), connecting these changes in CSH to Ca^2+^-mediated aggregation and our observed increase in phagocytosis. The aggregates formed by overcoming hydrophobic repulsion via Ca^2+^ interactions are phenotypically distinct from characterized cryptococcal biofilms with regards to phagocytosis by macrophages. These observations support the idea that the surface properties of the capsule are dynamic and play a critical role in both cell survival and the outcome of host infection (46). Further, biomineralization in other microbes suggest that calcium-phosphate microcrystals may form on the surface of the cryptococcal aggregates, though what role these microcrystals play in phagocytosis by macrophages remains to be seen, our data suggests they act as opsonins by reducing the anti-phagocytic properties of the capsule. These novel calcium-mediated cryptococcal aggregates suggest that the dication interactions are necessary to form the capsule structure and warrant further investigation, as they may be exploited as a therapeutic target for *Cryptococcus neoformans* infections.

### Experimental Procedures

#### Yeast strains, culture conditions, media, and solutions

*C. neoformans* serotype A reference strain H99, *cap59* deletion mutant (acapsular), single motif (motif 2) Mu-1 and ATCC 24064 (24064) were stored in -80 °C in glycerol stocks from our laboratory were used in preliminary aggregation assays. *C. neoformans* from frozen stocks was grown in Yeast Extract Peptone Dextrose (YPD) (BD DIFCO™ - Franklin Lakes, NJ, US) and Minimal Media (MM) (15 mM D-glucose, 10 mM MgSO_4_ x•7H_2_O, 20.3 mM KH_2_PO_4_, 3 mM glycine, 10 mg/mL thiamine, pH 5.5). Liquid cultures for aggregation assays and microscopy were grown at 30 °C with rotation at 150 rpm for 48 h. For capsular polysaccharide isolation, cells were grown for 48 h in YPD and subcultured into MM for 72 h. CaCl_2_ (Sigma), MgCl_2_ · 6H_2_O (Sigma), SrCl_2_· 6H_2_O (Chem Ipex), CdCl_2_ (Chem Ipex) and D-Glucuronic acid (Selleck Chemical) were analytical grade and prepared in deionized (Millipore, Milli-Q) water. Cells were either washed in Dulbecco’s phosphate buffered solution without Ca^2+^ or Mg^2+^ (Cytiva) or buffered with a 1 mM (1X) solution of phosphate diluted from a 10X preparation (8.5 mM Na_2_HPO_4_ and 1.5 mM KH_2_PO_4_, pH adjusted to 7.4).

#### Cellular Aggregation

*Cryptococcus neoformans* strain ATCC 24064 (serotype A) was grown in yeast peptone dextrose broth at 30 °C for 2 d, washed 3x and resuspended in DPBS. In a 24 well plate, cells were incubated with 10 mM of MgCl_2_, CaCl_2_, SrCl_2_, or CdCl_2_ and observed via light microscopy at 10x.

#### Capsular polysaccharide isolation

CPS was isolated as previously reported (47). Briefly, *C. neoformans* single motif strain 24064 was cultured in MM and washed 3x in DPBS without no Ca^2+^ or Mg^2+^. Cells were brought to a density of 1 x 10^8^ cells/ml. After washing, a suspension of cells in 2 ml of PBS was sonicated on ice using a horn sonicator (Fisher Scientific Sonic Dismembrator F550 W/ultrasonic Converter) on setting 3 for 30 s (8 W). Following sonication, cells were pelleted at 4,000 rpm and the supernatant (containing the sonicated capsule polysaccharide) was filter sterilized using a 0.22 µm filter. Capsule contained within the supernatant was quantified using size-exclusion chromatography multi-angle light scattering to determine size distribution of polymers and approximated molecular weight.

#### High Performance Liquid Chromatography and Multi-Angle Light Scattering

Capsular polysaccharide (CPS) was prepared as previously discussed. CPS samples were additionally prepared with the addition of 5 mM CaCl_2_ or MgCl_2_ with or without 1 mM phosphate. CPS with and without 1 mM phosphate were compared. Using a Shodex (Wiesbaden, Germay) SB-806M HQ OHPak column (8.0 mm ID x 300 mm) equipped with the OHPak SB-G guard column, CPS was quantified, and an estimate of molecular weight was determined by Size Exclusion Chromatography (SEC) on an Agilent (Santa Clara, California) 1200 HPLC coupled to a Wyatt (Santa Barbara, California) miniDawn Multi-Angle light scattering (MALS). The flow rate was 0.5 mL/min for this column. Analysis of sample retention time and peak isolation of CPS were analyzed with an 80-minute isocratic run, with HPLC-grade water as the mobile phase. The column temperature was 25 °C, sample injection volume was 5 µl for peak determination and 20 µl for peak collection. Flow rate was 0.2 mL/min on an Acclaim™ SEC-300 column (300 mm x 4.6 mm, Thermo Fisher Scientific). Both Shodex SB-806M HQ OHPak and Acclaim SEC-300 columns were standardized and calibrated using pullulan standards p5 (Mw: 5,000) and p50 (Mw: 50,000) from the Shodex Standard p-82 pullulan kit, using the Wyatt standardization protocol for the miniDawn MALS.

#### Dynamic Light Scattering

Capsule polysaccharide particle size distribution was determined with a Zeta Potential Analyzer (Brookhaven Instruments). A 100 µL sample of CPS was placed in a cuvette (Eppendorf) at room temperature. Data are expressed as the average of 10 runs of 1-minute data collection. The multimodal size distributions of the particles were obtained by a non-negatively constrained least squares algorithm based on the intensity of light scattered by each particle. The multimodal size distributions of particles from samples were graphed for comparison.

#### Nuclear Magnetic Resonance Experiments

Monosaccharide samples of glucuronic acid, mannose, and xylose were prepared in D_2_O at 50 and 100 mM and pH adjusted to 5.5. Metal (CaCl_2_, MgCl_2_, SrCl_2_, CdCl_2_) and phosphate solutions were prepared in D_2_O at 50 and 100 mM and added to monosaccharide samples immediately prior to spectral collection. All samples included D_6_-DSS as an internal standard. Both 1D ^1^H and Diffusion Order Spectroscopy experiments were collected on a Bruker Avance (500 MHz) spectrometer equipped with an XYZ gradient TCl cryoprobe. The work was done at the Food and Drug Administration, Laboratory of Bacterial Polysaccharides. DOSY data were analyzed using MNOVA (courtesy of NMRbox).

#### Mass Spectrometry

Samples collected from HPLC separation were concentrated by roughly 50% using a speed vacuum centrifuge. Samples were prepared by mixing 1 µL of 2,5-dihydroxybenzoic acid matrix (Super-DHB) (50% acetonitrile, 50% water, Sigma-Aldrich) with 1 µL of CPS sample, dotted onto a 96-spot stell target plate and allowed to dry. Samples were analyzed on the Voyager De-STR Matrix-assisted laser desorption/ionization time-of-flight (MALDI-TOF) mass spectrometry in linear mode to determine the mass of polysaccharide(s). Polymers were assigned based on the M2 motif and components of GXM, dication and phosphate additions, along with sodium or potassium adducts expected in MALDI-TOF analysis of polysaccharides.

#### Circular dichroism (CD)

CD measurements of CPS were carried out on a Jasco J-810 spectropolarimeter with a 2 mm cell at 25 °C with nitrogen flow at 22 (L/min). Samples contained 30 ug/mL of strain 24064 CPS dissolved in ultrapure water. A subset of CPS samples was treated with CaCl_2_ or MgCl_2_ with and without 1 mM phosphate. CD spectra were recorded between 185 and 350 nm and were expressed as molar ellipticity ([*θ*]), represented as mdeg/mv.

#### Scanning electron microscopy

A stationary phase culture of *C. neoformans* strain ATCC 24064 was washed and resuspended phosphate buffered saline treated (control) or with either 10 mM CaCl_2_ or MgCl_2_. Samples were fixed in 2.5% glutaraldehyde, 3 mM MgCl_2_, in 0.05 M sodium cacodylate buffer, pH 7.2 overnight at 4 °C. After buffer rinse, samples post fixed in 1% osmium tetroxide in 0.075 M sodium cacodylate buffer on ice in the dark for 1 h. Following a distilled water rinse, samples were dehydrated in a graded series of ethanol and left to dry overnight in a desiccator with hexamethyldisilazane (HMDS). Samples were mounted on carbon-coated stubs and imaged on JEOL Field Emission SEM (JSM-IT700HR InTouchScope) at 2kV. The work was done at the materials and characterization core at JHU (MCP).

#### Phagocytic Index assay

Phagocytosis assays were performed as previously described with minor modifications (48). BMDM were seeded (5 x 10^4^ cells/well) onto clear bottom 96 well plates. Cells were activated with IFN-gamma (10 ng/mL, Roche) and LPS (0.5 µg/mL) 18 h prior to *C. neoformans* infection and incubated at 37 °C with 9% CO_2_ overnight. On the following day, BMDMs were infected with *C. neoformans* strain 24064 at 5 x 10^4^ for an MOI of 1. *C. neoformans* was either opsonized with murine monoclonal antibody (mAb) IgG1 18B7 (10 µg/mL) or left unopsonized as a control. In parallel, *C. neoformans* was treated with 5 mM of CaCl_2_ and 1 mM of phosphate, incubated for 10 min, and washed 2X in MQ water. *C. neoformans* treated with CaCl_2_ and phosphate were similarly left unopsonized (control) and opsonized with 18B7 (10 µg/mL) prior to BMDM infection. BMDMs and *C. neoformans* were left to co-incubate at 37 °C, 9% CO_2_ for 2 h. Afterwards, media was removed, gently washed with fresh, pre-warmed, DMEM and stained with Uvitex 2B (Polysciences, Warrington, PA) for 10 min, to identify extracellular *C. neoformans*. BMDMs were again washed 2x with fresh medium. Images were taken using Agilent CYTATION 7 plate reader and imager with 20X objective. The phagocytic index was determined by the number of internalized cryptococcal cells per macrophage.

#### Immunofluorescence

Fluorescent staining of *C. neoformans*. Cells grown in YPD for 48 h were washed 3x and stained with Uvitex 2B. After washing twice in MQ water, cells were stained with 18B7 mAb direct-conjugated to Oregon green fluorophore (18B7-OG) for 1 h at 37 °C protected from light. Subsets of cells were either treated with 5 mM CaCl_2_ and 1 mM phosphate before or after 18B7-OG staining. In either case, after calcium-phosphate treatment, cells were washed 2x in MQ water.

#### ATR-FTIR Spectroscopy

Mid-infrared spectra (400 – 4000 cm^-1^) were collected on Thermo Scientific Nicolet IS5 FT-IR Spectrometer equipped with an iD5 ATR horizontal sampling accessory. The internal reflection element was a single-bounce diamond laminate. Before each experiment, the diamond was washed with ethanol to remove previous material and any debris. Spectra were acquired with 20 microliters of sample placed on top of the crystal. The spectral resolution for all experiments was 4 cm^-1^. A total of 32 interferograms were acquired and averaged for each spectrum.

#### Density Functional Theory (DFT) Computational Calculations

All calculations were carried out using Gaussian 16, rev. C02, with pre- and post-processing conducted in GaussView 6 (49, 50). Density Functional Theory (DFT) was employed with a meta-generalized gradient approximation as given by Tao, Perdew, Staroverov, and Scuseria (TPSS) (51). The Stuttgart, Dresden, Dusseldorf (SDD) effective core potential (ECP) basis set was employed for all atoms. This places 10 core electrons of Ca, 28 of Sr, and 28 of Cd into the modified Wood-Boring (MWB) electronic core potential (ECP) (52, 53). All other atoms utilized the Dunning-Huzinga valence double-zeta basis (54, 49). For the studies, the glucuronic acid residues only represent the D isomer, which is biologically relevant. Solvation was accounted for implicitly using the Integral Equation Formalism Polarizable Continuum Model (IEFPCM) (55). All stationary points were optimized without geometry or symmetry constraints with the Berny algorithm and confirmed with harmonic vibrational frequency analysis (56). The frequencies were calculated in solution with the harmonic oscillator approximations to confirm local minima and determine thermodynamic properties. The entropy of the computed structures was evaluated by employing the gas-phase rigid rotor and harmonic oscillator approximation. No hindered rotation correction was applied. To map the gas-phase potential energy surface (PES) Wolfram Language was used to generate an ensemble of 700 random conformers for each monosaccharide. The geometry of each conformer was optimized using the Merck Molecular Force Field (MMFF) and then the energy was calculated.

## Data Availability

Data is available upon request.

## Supporting Information

This article contains supporting information.

## Acknowledgements

We thank Daron Freedberg and Audra Hargett from the Laboratory of Bacterial Polysaccharides at the Food and Drug Administration for their invaluable assistance with collection and interpretation of the DOSY-NMR interactions between monosaccharides and metal dications, along with their feedback and comments on thus Manuscript. We are grateful to the Hardwick lab for providing laboratory space for sonication experiments and to the JHU Chemistry Department for providing us with a Thermo Scientific Nicolet IS5 FT-IR Spectrometer. We thank Barbara Smith of the Johns Hopkins Microscope Core Facility for expert assistance with SEM imaging. We also thank Dr. Robert Nachbar for his detailed review of the manuscript.

## Author contributions

G.R.G., J. L. S., M. P. W., and A. C. conceptualization; G.R.G., J. L. S., M. P. W., and A. C. formal analysis; A. C. funding acquisition; G. R. G. and M. P. W. investigation; G.R.G., J. L. S., M. P. W., and A. C. methodology; G.R.G., J. L. S., M. P. W., and A. C. project administration; G.R.G., J. L. S., M. P. W., and A. C. supervision; G.R.G., J. L. S., M. P. W., and A. C. validation; G. R. G. and A. C. visualization; G. R. G. writing—original draft; G.R.G., J. L. S., M. P. W., S. R., A. H., D. I. F., and A. C. writing—review and editing.

## Funding and Additional Information

This work was supported by the National Institutes of Health, grant R01AI152078. The content is solely the responsibility of the authors and does not necessarily represent the official views of the National Institutes of Health.

## References

1. Rajasingham, R., Smith, R. M., Park, B. J., Jarvis, J. N., Govender, N. P., Chiller, T. M., Denning, D. W., Loyse, A., and Boulware, D. R. (2017) Global burden of disease of HIV-associated cryptococcal meningitis: an updated analysis. Lancet Infect. Dis. 17, 873–881

2. Vu, K., Tham, R., Uhrig, J. P., Thompson, G. R., Na Pombejra, S., Jamklang, M., Bautos, J. M., and Gelli, A. (2014) Invasion of the central nervous system by Cryptococcus neoformans requires a secreted fungal metalloprotease. MBio. 5, e01101–14

3. Nosanchuk, J. D., and Casadevall, A. (1997) Cellular charge of Cryptococcus neoformans: contributions from the capsular polysaccharide, melanin, and monoclonal antibody binding. Infect. Immun. 65, 1836–1841

4. Nicola, A. M., Frases, S., and Casadevall, A. (2009) Lipophilic dye staining of Cryptococcus neoformans extracellular vesicles and capsule. Eukaryotic Cell. 8, 1373–1380

5. Bose, I., Reese, A. J., Ory, J. J., Janbon, G., and Doering, T. L. (2003) A yeast under cover: the capsule of Cryptococcus neoformans. Eukaryotic Cell. 2, 655–663

6. Casadevall, A., Coelho, C., Cordero, R. J. B., Dragotakes, Q., Jung, E., Vij, R., and Wear, M. P. (2019) The capsule of Cryptococcus neoformans. Virulence. 10, 822–831

7. Gates, M. A., Thorkildson, P., and Kozel, T. R. (2004) Molecular architecture of the Cryptococcus neoformans capsule. Mol. Microbiol. 52, 13–24

8. Cherniak, R., Reiss, E., Slodki, M. E., Plattner, R. D., and Blumer, S. O. (1980) Structure and antigenic activity of the capsular polysaccharide of Cryptococcus neoformans serotype A. Mol. Immunol. 17, 1025–1032

9. Merrifield, E. H., and Stephen, A. M. (1980) Structural investigations of two capsular polysaccharides from cryptococcus neoformans. Carbohydr. Res. 86, 69–76

10. McFadden, D., Zaragoza, O., and Casadevall, A. (2006) The capsular dynamics of Cryptococcus neoformans. Trends Microbiol. 14, 497–505

11. McFadden, D. C., De Jesus, M., and Casadevall, A. (2006) The physical properties of the capsular polysaccharides from Cryptococcus neoformans suggest features for capsule construction. J. Biol. Chem. 281, 1868–1875

12. Cherniak, R., Valafar, H., Morris, L. C., and Valafar, F. (1998) Cryptococcus neoformans chemotyping by quantitative analysis of 1H nuclear magnetic resonance spectra of glucuronoxylomannans with a computer-simulated artificial neural network. Clin. Diagn. Lab. Immunol. 5, 146–159

13. Kozel, T. R., Levitz, S. M., Dromer, F., Gates, M. A., Thorkildson, P., and Janbon, G. (2003) Antigenic and biological characteristics of mutant strains of Cryptococcus neoformans lacking capsular O acetylation or xylosyl side chains. Infect. Immun. 71, 2868–2875

14. Maxson, M. E., Dadachova, E., Casadevall, A., and Zaragoza, O. (2007) Radial mass density, charge, and epitope distribution in the Cryptococcus neoformans capsule. Eukaryotic Cell. 6, 95–109

15. Nimrichter, L., Frases, S., Cinelli, L. P., Viana, N. B., Nakouzi, A., Travassos, L. R., Casadevall, A., and Rodrigues, M. L. (2007) Self-aggregation of Cryptococcus neoformans capsular glucuronoxylomannan is dependent on divalent cations. Eukaryotic Cell. 6, 1400–1410

16. Frases, S., Pontes, B., Nimrichter, L., Rodrigues, M. L., Viana, N. B., and Casadevall, A. (2009) The elastic properties of the Cryptococcus neoformans capsule. Biophys. J. 97, 937–945

17. Robertson, E. J., Wolf, J. M., and Casadevall, A. (2012) EDTA inhibits biofilm formation, extracellular vesicular secretion, and shedding of the capsular polysaccharide glucuronoxylomannan by Cryptococcus neoformans. Appl. Environ. Microbiol. 78, 7977–7984

18. Rathore, S. S., Raman, T., and Ramakrishnan, J. (2016) Magnesium Ion Acts as a Signal for Capsule Induction in Cryptococcus neoformans. Front. Microbiol. 7, 325

19. Rivera, J., Feldmesser, M., Cammer, M., and Casadevall, A. (1998) Organ-dependent variation of capsule thickness in Cryptococcus neoformans during experimental murine infection. Infect. Immun. 66, 5027–5030

20. Vecchiarelli, A., Pericolini, E., Gabrielli, E., Kenno, S., Perito, S., Cenci, E., and Monari, C. (2013) Elucidating the immunological function of the Cryptococcus neoformans capsule. Future Microbiol. 8, 1107–1116

21. Danchik, C., and Casadevall, A. (2020) Role of Cell Surface Hydrophobicity in the Pathogenesis of Medically-Significant Fungi. Front. Cell. Infect. Microbiol. 10, 594973

22. Patergnani, S., Danese, A., Bouhamida, E., Aguiari, G., Previati, M., Pinton, P., and Giorgi, C. (2020) Various aspects of calcium signaling in the regulation of apoptosis, autophagy, cell proliferation, and cancer. Int. J. Mol. Sci. 10.3390/ijms21218323

23. Jastrzab, R., Nowak, M., Skrobanska, M., and Zabiszak, M. (2016) Complexation copper(II) or magnesium ions with -glucuronic acid – potentiometric, spectral and theoretical studies. J. Coord. Chem. 69, 2174–2181

24. Gagné, O. C., and Hawthorne, F. C. (2016) Bond-length distributions for ions bonded to oxygen: alkali and alkaline-earth metals. Acta Crystallogr. B Struct. Sci. Cryst. Eng. Mater. 72, 602–625

25. Pauling, L. (1940) The Nature of the Chemical Bond and the Structure of Molecules and Crystals; an introduction to modern structural chemistry, second edition, Cornell University Press

26. Casadevall, A., Cleare, W., Feldmesser, M., Glatman-Freedman, A., Goldman, D. L., Kozel, T. R., Lendvai, N., Mukherjee, J., Pirofski, L. A., Rivera, J., Rosas, A. L., Scharff, M. D., Valadon, P., Westin, K., and Zhong, Z. (1998) Characterization of a murine monoclonal antibody to Cryptococcus neoformans polysaccharide that is a candidate for human therapeutic studies. Antimicrob. Agents Chemother. 42, 1437–1446

27. De Leon-Rodriguez, C. M., Fu, M. S., Çorbali, M. O., Cordero, R. J. B., and Casadevall, A. (2018) The Capsule of Cryptococcus neoformans Modulates Phagosomal pH through Its Acid-Base Properties. mSphere. 10.1128/mSphere.00437-18

28. Ueda, Y., and Taira, Z. (2013) Effect of anions or foods on absolute bioavailability of calcium from calcium salts in mice by pharmacokinetics. J. Exp. Pharmacol. 5, 65–71

29. Fernández, I., Holzmann, N., and Frenking, G. (2020) The Valence Orbitals of the Alkaline-Earth Atoms. Chem. Eur. J. 26, 14194–14210

30. Pearson, R. G. (1963) Hard and Soft Acids and Bases. J. Am. Chem. Soc. 85, 3533–3539

31. Fu, Y., Huang, X., and Zhou, Z. (2020) Insight into the Assembling Mechanism of Cryptococcus Capsular Glucuronoxylomannan Based on Molecular Dynamics Simulations. ACS Omega. 5, 29351–29356

32. Kovács, A., Nemcsok, D. S., and Kocsis, T. (2010) Bonding interactions in EDTA complexes. Journal of Molecular Structure: THEOCHEM. 950, 93–97

33. Sobeck, D. C., and Higgins, M. J. (2002) Examination of three theories for mechanisms of cation-induced bioflocculation. Water Res. 36, 527–538

34. Agmo Hernández, V. (2023) An overview of surface forces and the DLVO theory. ChemTexts. 9, 10

35. Arias, J. L., and Fernández, M. S. (2008) Polysaccharides and proteoglycans in calcium carbonate-based biomineralization. Chem. Rev. 108, 4475–4482

36. Lee, H. H., Del Pozzo, J., Salamanca, S. A., Hernandez, H., and Martinez, L. R. (2019) Reduced phagocytosis and killing of Cryptococcus neoformans biofilm-derived cells by J774.16 macrophages is associated with fungal capsular production and surface modification. Fungal Genet. Biol. 132, 103258

37. Pruitt, H. M., Zhu, J. C., Riley, S. P., and Shi, M. (2025) The Hidden Fortress: A Comprehensive Review of Fungal Biofilms with Emphasis on Cryptococcus neoformans. Journal of Fungi. 11, 236

38. Frankel, R. B. (2003) Biologically induced mineralization by bacteria. Reviews in Mineralogy and Geochemistry. 54, 95–114

39. Fishman, M. R., Giglio, K., Fay, D., and Filiatrault, M. J. (2018) Physiological and genetic characterization of calcium phosphate precipitation by Pseudomonas species. Sci. Rep. 8, 10156

40. Li, Q., Csetenyi, L., and Gadd, G. M. (2014) Biomineralization of metal carbonates by Neurospora crassa. Environ. Sci. Technol. 48, 14409–14416

41. Li, Q., Csetenyi, L., Paton, G. I., and Gadd, G. M. (2015) CaCO3 and SrCO3 bioprecipitation by fungi isolated from calcareous soil. Environ. Microbiol. 17, 3082–3097

42. Fu, M. S., Coelho, C., De Leon-Rodriguez, C. M., Rossi, D. C. P., Camacho, E., Jung, E. H., Kulkarni, M., and Casadevall, A. (2018) Cryptococcus neoformans urease affects the outcome of intracellular pathogenesis by modulating phagolysosomal pH. PLoS Pathog. 14, e1007144

43. Baker, R. P., and Casadevall, A. (2023) Reciprocal modulation of ammonia and melanin production has implications for cryptococcal virulence. Nat. Commun. 14, 849

44. Vij, R., Danchik, C., Crawford, C., Dragotakes, Q., and Casadevall, A. (2020) Variation in Cell Surface Hydrophobicity among Cryptococcus neoformans Strains Influences Interactions with Amoebas. mSphere. 10.1128/mSphere.00310-20

45. Radosa, S., and Hillmann, F. (2021) Host-pathogen interactions: lessons from phagocytic predation on fungi. Curr. Opin. Microbiol. 62, 38–44

46. Li, L., Zaragoza, O., Casadevall, A., and Fries, B. C. (2006) Characterization of a flocculation-like phenotype in Cryptococcus neoformans and its effects on pathogenesis. Cell. Microbiol. 8, 1730–1739

47. Wear, M. P., Jacobs, E., Wang, S., McConnell, S. A., Bowen, A., Strother, C., Cordero, R. J. B., Crawford, C. J., and Casadevall, A. (2022) Cryptococcus neoformans capsule regrowth experiments reveal dynamics of enlargement and architecture. J. Biol. Chem. 298, 101769

48. Nicola, A. M., and Casadevall, A. (2012) In vitro measurement of phagocytosis and killing of Cryptococcus neoformans by macrophages. Methods Mol. Biol. 844, 189–197

49. Frisch, M. J., Trucks, G. W., Schlegel, H. B., Scuseria, G. E., Robb, M. A., Cheeseman, J. R., Scalmani, G., Barone, V., Petersson, G. A., Nakatsuji, H., Li, X., Caricato, M., Marenich, A. V., Bloino, J., Janesko, B. G., Gomperts, R., Mennucci, B., Hratchian, H. P., Ortiz, J. V., Izmaylov, A. F., Sonnenberg, J. L., Williams, Ding, F., Lipparini, F., Egidi, F., Goings, J., Peng, B., Petrone, A., Henderson, T., Ranasinghe, D., Zakrzewski, V. G., Gao, J., Rega, N., Zheng, G., Liang, W., Hada, M., Ehara, M., Toyota, K., Fukuda, R., Hasegawa, J., Ishida, M., Nakajima, T., Honda, Y., Kitao, O., Nakai, H., Vreven, T., Throssell, K., Montgomery Jr., J. A., Peralta, J. E., Ogliaro, F., Bearpark, M. J., Heyd, J. J., Brothers, E. N., Kudin, K. N., Staroverov, V. N., Keith, T. A., Kobayashi, R., Normand, J., Raghavachari, K., Rendell, A. P., Burant, J. C., Iyengar, S. S., Tomasi, J., Cossi, M., Millam, J. M., Klene, M., Adamo, C., Cammi, R., Ochterski, J. W., Martin, R. L., Morokuma, K., Farkas, O., Foresman, J. B., and Fox, D. J. (2016) Gaussian 16 Rev. *C.02*, Wallingford, CT

50. Dennington, R., Keith, T. A., and Milliam, J. M. (2016) GaussView, Semichem Incorporated, Shawnee Mission, KS

51. Staroverov, V. N., Scuseria, G. E., Tao, J., and Perdew, J. P. (2003) Comparative assessment of a new nonempirical density functional: Molecules and hydrogen-bonded complexes. J. Chem. Phys. 119, 12129–12137

52. Kaupp, M., Schleyer, P. v. R., Stoll, H., and Preuss, H. (1991) Pseudopotential approaches to Ca, Sr, and Ba hydrides. Why are some alkaline earth MX2 compounds bent? J. Chem. Phys. 94, 1360–1366

53. Andrae, D., Huermann, U., Dolg, M., Stoll, H., and Preu, H. (1990) Energy-adjustedab initio pseudopotentials for the second and third row transition elements. Theor. Chim. Acta. 77, 123–141

54. Dunning, Thom. H., and Hay, P. J. (1977) Gaussian basis sets for molecular calculations. in Methods of electronic structure theory (Schaefer, H. F. ed), pp. 1–27, Springer US, Boston, MA, 10.1007/978-1-4757-0887-5_1

55. Tomasi, J., Mennucci, B., and Cammi, R. (2005) Quantum mechanical continuum solvation models. Chem. Rev. 105, 2999–3093

56. Schlegel, H. B. (1982) Optimization of equilibrium geometries and transition structures. J. Comput. Chem. 3, 214–218

