## Supplemental Materials for "Calcium triggers *Cryptococcus neoformans* aggregation by forming coordination bonds with capsular glucuronoxylomannan"

Supporting Information

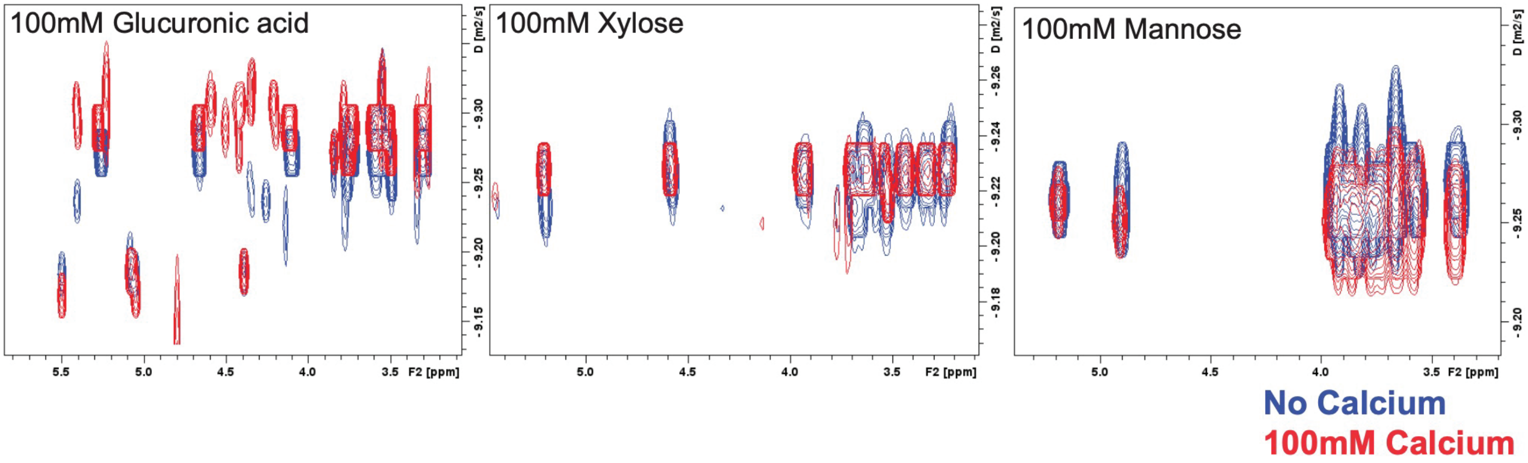


**Supporting Figure 1.** Diffusion NMR experiments to examine interaction between GXM monosaccharide residues and calcium cations. [$^1$H] DOSY spectra of A. D-Glucuronic acid B. Xylose or C. Mannose in 1x PBS with (red) and without (blue) 50 mM calcium chloride added.

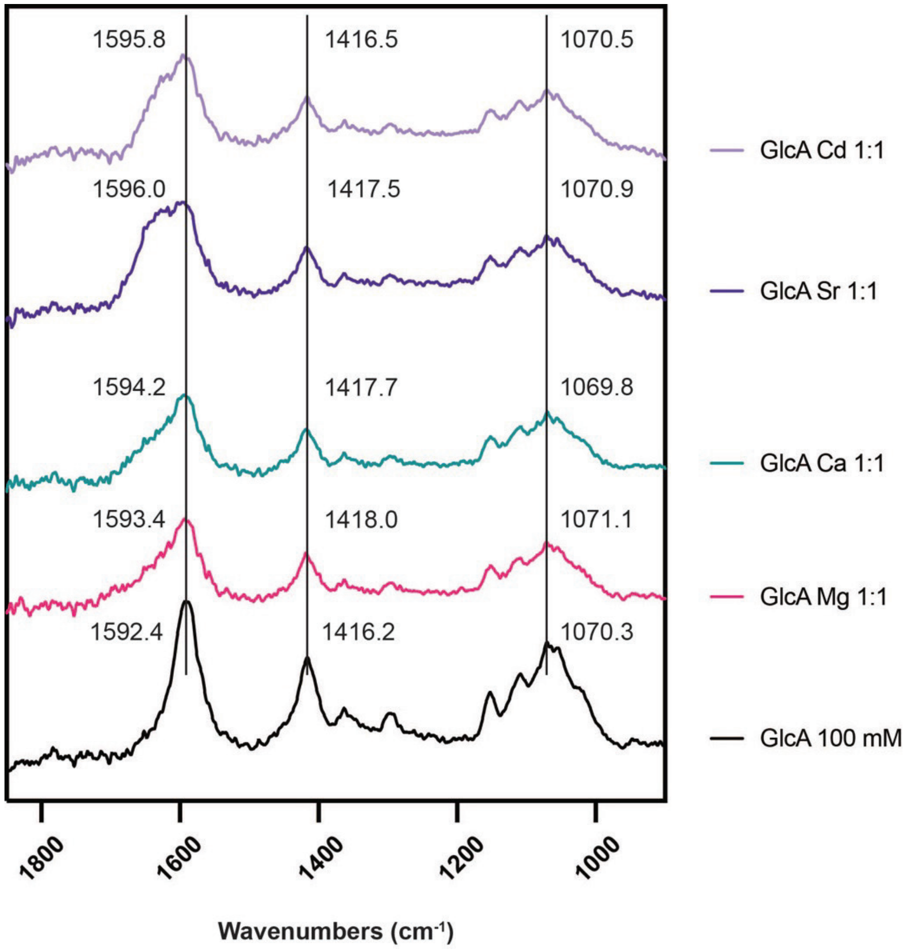


**Supporting Figure 2**: Attenuated Total Reflectance-Fourier Transform infrared spectroscopy (ATR-FTIR) of GlcA in 1xPBS (black) and with 1:1 molar ratio of $CaCl_2$ (teal), $MgCl_2$ (pink), $SrCl_2$ (purple) or $CdCl_2$ (lilac).

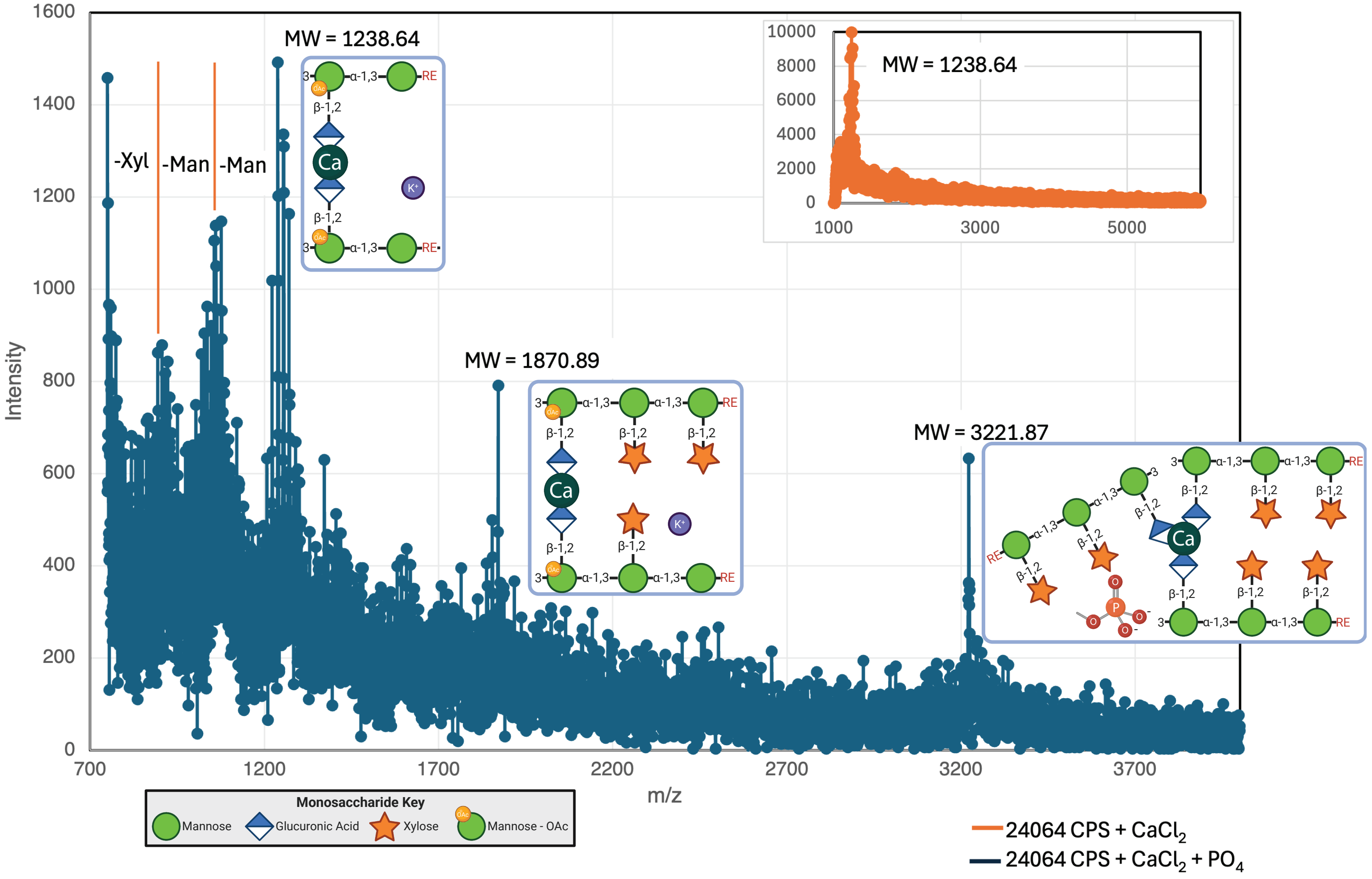


**Suppor ing Figure 3:** Matrix Assisted aser Desorption Ioni ation - Time of Flight (MALDI-TOF) Mass Spectrometry analysis of Cryptococcal CPS with added calcium. Peak m analysis using molecular weights compared to GXM mo f polymer masses considering calcium coordina on and MS adducts.

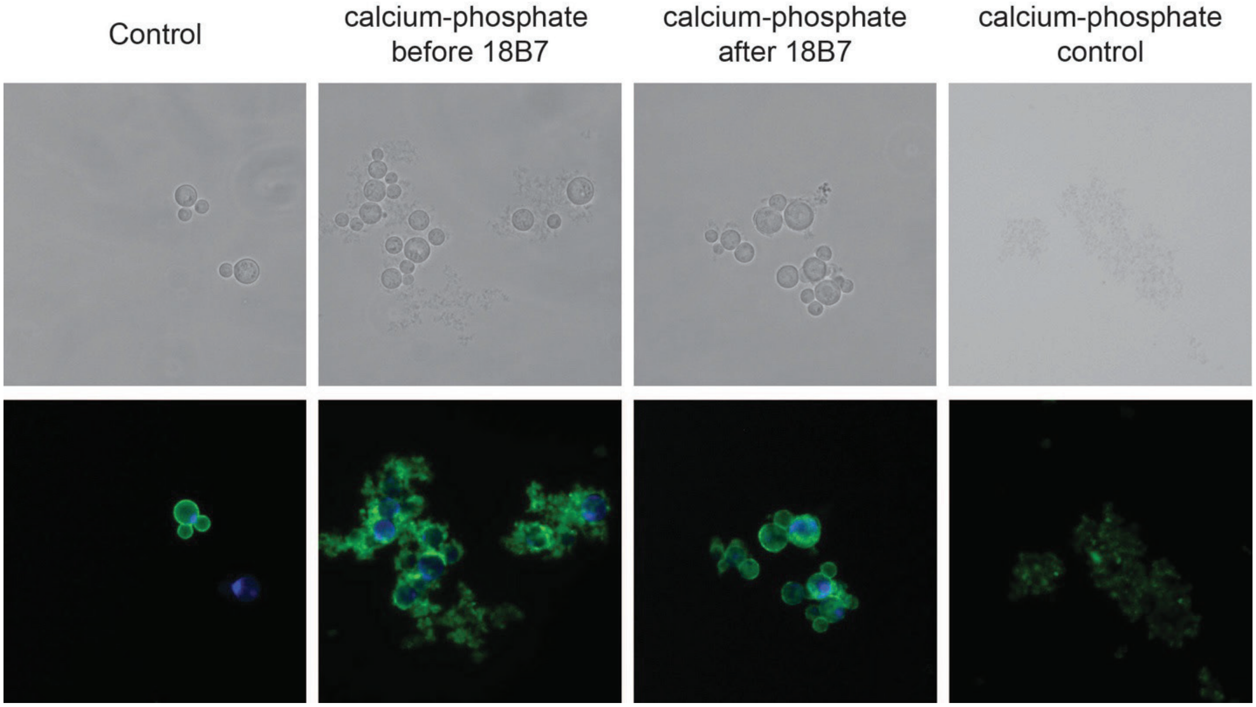


**Supporting Figure 4:** Immunofluorescence microscopy of *C. neoformans* cells stained with fluorescently labeled monoclonal antibody 18B7. Cells imaged at 40x. Top panel: Bright Field, Bottom panel: Green.

**Supporting Table 1**: Calculated electronic and zero-point energies in bulk solvent

| Compound | Coordination site | GlcA:$M^{2+}$ ratio | E(elec) (Hartree) | E(ZPE) (Hartree) | E(elec) + E(ZPE) |
|---|---|---|---|---|---|
| $Ca^{2+}$ | NA | Dication calcium | -36.664152 | 0 | -36.664152 |
| $Mg^{2+}$ | NA | Dication magnesium | -199.86662 | 0 | -199.86662 |
| $Sr^{2+}$ | NA | Dication strontium | -30.585463 | 0 | -30.585463 |
| $Cd^{2+}$ | NA | Dication cadmium | -167.39026 | 0 | -167.39026 |
| $C_6H_{10}O_7$ | NA | Neutral GlcA | -761.25117 | 0.170977 | -761.08019 |
| $[C_6H_9O_7]^-$ | NA | Anion GlcA | -760.80167 | 0.158234 | -760.64344 |
| $[C_6H_9O_7]Ca^+$ | Metal to $COO^-$ | 1GlcA:1Ca | -797.50748 | 0.160891 | -797.34659 |
| $[C_6H_9O_7]Mg^+$ | | 1GlcA:1Mg | -960.72109 | 0.161428 | -960.55966 |
| $[C_6H_9O_7]Sr^+$ | | 1GlcA:1Sr | -791.42372 | 0.160463 | -791.26325 |
| $[C_6H_9O_7]Cd^+$ | | 1GlcA:1Cd | -928.23657 | 0.160791 | -928.07578 |
| $[2C_6H_9O_7]Ca$ | | 2GlcA:1Ca | -1558.3475 | 0.321188 | -1558.0263 |
| $[2C_6H_9O_7]Mg$ | | 2GlcA:1Mg | -1721.5734 | 0.322973 | -1721.2504 |
| $[2C_6H_9O_7]Sr$ | | 2GlcA:1Sr | -1552.258 | 0.320552 | -1551.9375 |
| $[2C_6H_9O_7]Cd$ | | 2GlcA:1Cd | -1689.081 | 0.321855 | -1688.7591 |
| $[3C_6H_9O_7]Ca^-$ | | 3GlcA:1Ca | -2319.1828 | 0.480831 | -2318.702 |
| $[3C_6H_9O_7]Mg^-$ | | 3GlcA:1Mg | -2482.4184 | 0.483141 | -2481.9352 |
| $[3C_6H_9O_7]Sr^-$ | | 3GlcA:1Sr | -2313.0911 | 0.480394 | -2312.6107 |
| $[3C_6H_9O_7]Cd^-$ | | 3GlcA:1Cd | -2449.909 | 0.481669 | -2449.4274 |
| $[4C_6H_9O_7]Ca^{2-}$ | | 4GlcA:1Ca | -3080.0121 | 0.641704 | -3079.3704 |
| $[4C_6H_9O_7]Mg^{2-}$ | | 4GlcA:1Mg | -3243.2546 | 0.64227 | -3242.6123 |
| $[4C_6H_9O_7]Sr^{2-}$ | | 4GlcA:1Sr | -3073.9132 | 0.639773 | -3073.2735 |
| $[4C_6H_9O_7]Cd^{2-}$ | | 4GlcA:1Cd | -3210.7459 | 0.644416 | -3210.1015 |
| $[C_6H_9O_7]Ca^+$ | Metal to ring | 1GlcA:1Ca | -797.52008 | 0.161279 | -797.3588 |
| $[C_6H_9O_7]Mg^+$ | | 1GlcA:1Mg | -960.7379 | 0.162373 | -960.57553 |
| $[C_6H_9O_7]Sr^+$ | | 1GlcA:1Sr | -791.43368 | 0.160904 | -791.27278 |
| $[C_6H_9O_7]Cd^+$ | | 1GlcA:1Cd | -928.24161 | 0.161098 | -928.08051 |
| $[2C_6H_9O_7]Ca$ | | 2GlcA:1Ca | -1558.3782 | 0.322608 | -1558.0556 |
| $[2C_6H_9O_7]Mg$ | | 2GlcA:1Mg | -1721.6182 | 0.325089 | -1721.2931 |
| $[2C_6H_9O_7]Sr$ | | 2GlcA:1Sr | -1552.2869 | 0.322178 | -1551.9647 |
| $[2C_6H_9O_7]Cd$ | | 2GlcA:1Cd | -1689.0963 | 0.322775 | -1688.7736 |
| $[3C_6H_9O_7]Ca^-$ | | 3GlcA:1Ca | -2319.212 | 0.483271 | -2318.7288 |
| $[3C_6H_9O_7]Mg^-$ | | 3GlcA:1Mg | -2482.4493 | 0.486079 | -2481.9632 |
| $[3C_6H_9O_7]Sr^-$ | | 3GlcA:1Sr | -2313.1148 | 0.481773 | -2312.6331 |
| $[3C_6H_9O_7]Cd^-$ | | 3GlcA:1Cd | -2449.9186 | 0.483772 | -2449.4348 |
| $C_6H_{10}O_7$ | Metal to hydroxyl | neutral GlcA* | -761.2578 | 0.171353 | -761.08644 |
| $[C_6H_9O_7]^-$ | | anion GlcA* | -760.79369 | 0.158634 | -760.63506 |
| $[C_6H_9O_7]Ca^+$ | | 1GlcA:1Ca | -797.51834 | 0.161586 | -797.35676 |
| $[C_6H_9O_7]Mg^+$ | | 1GlcA:1Mg | -960.73874 | 0.162758 | -960.57598 |
| $[C_6H_9O_7]Sr^+$ | | 1GlcA:1Sr | -791.42282 | 0.161879 | -791.26094 |
| $[C_6H_9O_7]Cd^+$ | | 1GlcA:1Cd | -928.23621 | 0.161463 | -928.07475 |
| $[2C_6H_9O_7]Ca$ | | 2GlcA:1Ca | -1558.3633 | 0.322661 | -1558.0406 |
| $[2C_6H_9O_7]Mg$ | | 2GlcA:1Mg | -1721.606 | 0.325174 | -1721.2808 |
| $[2C_6H_9O_7]Sr$ | | 2GlcA:1Sr | -1552.2671 | 0.32199 | -1551.9451 |
| $[2C_6H_9O_7]Cd$ | | 2GlcA:1Cd | -1689.0776 | 0.323147 | -1688.7544 |
| $[3C_6H_9O_7]Ca^-$ | | 3GlcA:1Ca | -2319.1994 | 0.483351 | -2318.7161 |
| $[3C_6H_9O_7]Mg^-$ | | 3GlcA:1Mg | -2482.4493 | 0.486366 | -2481.9629 |
| $[3C_6H_9O_7]Sr^-$ | | 3GlcA:1Sr | -2313.0982 | 0.482246 | -2312.616 |
| $[3C_6H_9O_7]Cd^-$ | | 3GlcA:1Cd | -2449.8997 | 0.483158 | -2449.4166 |
| $[4C_6H_9O_7]Ca^{2-}$ | | 4GlcA:1Ca | -3080.0229 | 0.644997 | -3079.3779 |
| $[4C_6H_9O_7]Mg^{2-}$ | | 4GlcA:1Mg | -3243.2661 | 0.647838 | -3242.6182 |
| $[4C_6H_9O_7]Sr^{2-}$ | | 4GlcA:1Sr | -3073.9161 | 0.64311 | -3073.273 |
| $[4C_6H_9O_7]Cd^{2-}$ | | 4GlcA:1Cd | -3210.7113 | 0.644199 | -3210.0671 |

*These structures were modified so that the hydroxyl groups were facing towards the ring in a pseudo ring configuration; only used for the metal:hydroxyl calculations.

**Supporting Table 2:** Δ of $\Delta_r H$ of reaction energies, $[M^{II} (D\text{-}GlcA)_n]^{2-n} + nD\text{-}GlcA^-$ ⍰$[M (D\text{-}GlcA)_{n+1}]^{2-n-1}$

| Binding reaction | Δ of $\Delta_r H$ | Mg | Ca | Sr | Cd |
|---|---|---|---|---|---|
| 1 | n=0 | -102.4 | -130.2 | -90.18 | -110.5 |
| | n=1 | -95.23 | -124.2 | -80.89 | -104.7 |
| | n=2 | -84.7 | -108.6 | -78.13 | -65.27 |
| | n=3 | -65.53 | -88.37 | -50.83 | -80.5 |
| 2 | n=0 | -134.5 | -171.9 | -115.2 | -122.9 |
| | n=1 | -140.1 | -194.6 | -127.3 | -130.4 |
| | n=2 | -78.14 | -70 | -65.53 | -46.63 |
| 3 | n=0 | -119.24 | -163.23 | -74.27 | -97.93 |
| | n=: | -96.22 | -151.31 | -97.06 | -85.22 |
| | n=2 | -74.33 | -91.66 | -62.25 | -39.41 |
| | n=3 | -38.36 | -21.29 | -25.76 | -8.69 |
